# Modulation of premotor cortex excitability mitigates the behavioral and electrophysiological abnormalities in a Parkinson’s Disease Mouse Model

**DOI:** 10.1101/2023.08.07.552205

**Authors:** In Sun Choi, Jinmo Kim, Joon Ho Choi, Ji-Woong Choi, Jong-Cheol Rah

## Abstract

The subthalamic nucleus (STN) is important in halting ongoing behaviors, referred to as stop-signal responses. The prefrontal regions innervating the STN exhibit increased activity during the stop-signal responses, and optogenetic activation of these neurons inhibits impulsive actions in rodents. High-frequency electrical stimulation of the STN effectively treats motor symptoms of Parkinson’s disease (PD), yet its underlying circuit mechanisms are unclear. Here, we investigated the involvement of STN-projecting premotor (M2) neurons in PD mouse models and the impact of deep brain stimulation targeting the STN (DBS-STN). We found that M2 neurons exhibited enhanced burst firing and synchronous oscillations in the PD mouse model. Remarkably, high-frequency stimulation of STN-projecting M2 neurons, simulating antidromic activation, during DBS-STN relieved motor symptoms and hyperexcitability. These changes were attributed to reduced firing frequency vs. current relationship through normalized hyperpolarization-activated inward current (I_h_). The M2 neurons in the PD model mouse displayed increased I_h_, which was reversed by high-frequency stimulation. Additionally, the infusion of ZD7288, an HCN channel blocker, into the M2 replicated the effects of high-frequency stimulation. In conclusion, our study unveils excessive excitability and suppressive motor control through M2-STN synapses in the PD mouse model. Antidromic excitation of M2 neurons during deep brain stimulation of the STN alleviates this suppression, thereby improving motor impairment. These findings provide insights into the circuit-level dynamics underlying deep brain stimulation’s therapeutic effects in PD. Targeting M2-STN synapses may be useful for future therapeutic strategies.

**Significance Statement:** Our study provides insight into the mechanisms underlying motor impairment in Parkinson’s disease (PD) and the therapeutic effect of deep brain stimulation (DBS). By elucidating the role of hyperpolarization-activated cyclic nucleotide-gated (HCN) channels in modulating the hyperexcitability of M2 neurons, we identify potential targets for future PD treatment strategies. Understanding the excessive excitability and suppressive motor control through M2-STN synapses in PD mouse models contributes to our knowledge of circuit-level dynamics underlying DBS effects. These findings have significant implications for improving motor symptoms and advancing the development of more precise and effective interventions for PD patients.

**Highlights:**

- STN-projecting M2 neurons show increased burst firing in PD mice.
- High-frequency stimulation relieves M2 hyperexcitability and motor symptoms.
- Reduction in M2 hyperexcitability by DBS linked to normalized I_h_.
- ZD7288 infusion in M2 replicates high-frequency stimulation effects.
- Targeted modulation of M2 neurons may alleviate PD symptoms.

## Introduction

The balance between movement and suppression of movement is finely regulated by corticobasal ganglia pathways (Aron and Poldrack, 2006; Schmidt et al., 2013), and disruption of the balance can lead to motor dysfunctions such as Parkinson’s disease (PD) or Huntington’s disease. The subthalamic nucleus (STN) transmits glutamatergic projections mainly to the substantia nigra pars reticulata (SNr) and the internal division of the globulus pallidum (GPi), both of which form gamma-aminobutyric acid (GABA)-ergic connections to the motor thalamus (Emmi et al., 2020). The activity of the STN can be predicted to suppress motor thalamocortical connections and, thus, action execution. Indeed, STN activity is involved in stopping or pausing planned or initiated movement across species. STN lesions shortened the reaction time in response to a go signal, increased the level of spontaneous locomotion, and impaired the ability to stop the ongoing movement (Eagle et al., 2008; Fife et al., 2017; Hamada and DeLong, 1992; Schmidt et al., 2013). Furthermore, functional removal of the STN improved motor symptoms in PD animal models, such as akinesia and rigidity (Aziz et al., 1991; Bergman et al., 1990; Whittier and Mettler, 1949). In line with these findings, increased firing of STN neurons has been observed in both PD patients and animal models (Bergman et al., 1994; Breit et al., 2007; Hassani et al., 1996; Magill et al., 2000; Remple et al., 2011).

The exact circuit mechanism of deep brain stimulation on the STN (DBS-STN), an efficient neurosurgical technique used to treat various neurological symptoms in PD, is still under debate (Lozano et al., 2019). However, one explanation is that depletion of the readily releasable pool (RRP) of neurotransmitters by high-frequency stimulation during DBS shuts down the abnormally exaggerated stopping effect of the STN (Lozano et al., 2019; Montgomery and Baker, 2000). Corroborating this idea, selective RRP depletion of STN axons leads to accelerated locomotor activity (Schweizer et al., 2014). However, the origin of the enhanced STN activity in PD remains unclear.

High-order motor cortical areas have been shown to control motor activity cessation in primates. The right inferior frontal gyrus of premotor cortex (Aron et al., 2003; Rubia et al., 2003) and pre-supplementary motor area (pre-SMA) (Li et al., 2006; Nachev et al., 2007; Swann et al., 2012) are reported to control motor inhibition via beta-frequency oscillations (Schaum et al., 2021; Sundby et al., 2021). Repetitive transcranial magnetic stimulation (rTMS) of these areas significantly weakened beta-frequency oscillation, reduced the ability to stop an action, and alleviated motor symptoms in patients (Mi et al., 2020, 2019; Sundby et al., 2021). Furthermore, relative hyperactivity in premotor cortex was observed in PD patients (Buhmann et al., 2004; Catalan et al., 1999; Samuel et al., 1997). Interestingly, these areas have strong direct synaptic connections to the STN (Aron et al., 2007a; Forstmann et al., 2010). Similarly, the rodent premotor cortex (M2), involved in motor planning and switching motor plans, has strong projections to the STN (Barthas and Kwan, 2017). During a task in which mice were trained to initiate or inhibit a motor action in response to an auditory cue, STN-projecting neurons in the deep layer of the prefrontal areas were preferentially active in no-go trials, and the tendency to initiate motor responses could be manipulated by the neurons (Kamigaki and Dan, 2017; Li et al., 2020). M2-STN projections appear to play a role in both stopping and planning to stop locomotive activity (Adam et al., 2022). Collectively, these observations suggest that overexcitation of high-order motor areas, leading to increased motor suppression via hyperdirect M2-STN pathway, may contribute to the etiology of PD. Furthermore, retrograde stimulation of M2 during DBS-STN could underlie some of the therapeutic effects observed. This idea was supported by the observation that optogenetic stimulation of M2 axons modulated the locomotor activity in dopamine-depleted mice (Magno et al., 2019) and by the therapeutic effect of rTMS on premotor areas (Mi et al., 2020, 2019; Sundby et al., 2021). However, while the effect of DBS-STN on the primary motor cortex (M1) has become clearer (Gradinaru et al., 2009; Johnson et al., 2015; Li et al., 2012a; Payoux et al., 2004; Sanders and Jaeger, 2016; Valverde et al., 2020), the role of high-frequency antidromic activation of M2 in these processes remains less understood.

Hyperpolarization-activated cyclic-nucleotide gated (HCN) channels, the cation channels activated upon hyperpolarization, exhibit dichotomous effects on circuit activity regulation. On one hand, HCN channels contribute to the depolarization of the resting membrane potential as HCN channels are active under -50 mV, thereby promoting action potential generation (Pape, 2003). Furthermore, hyperpolarization-induced rebound depolarization triggers subsequent action potentials in certain neurons (Chan et al., 2004a; Nolan et al., 2007; Robinson and Siegelbaum, 2003). Conversely, channel conductance in resting neurons reduces input resistance, leading to decreased integration and regenerative activity of dendrites (Huang et al., 2009; Magee, 1999). In addition to modulating neuronal excitability, HCN channels have been implicated in synchronizing neural networks to theta-frequency oscillation (Hu et al., 2009; Narayanan and Johnston, 2008; Pike et al., 2000; Rotstein et al., 2005; Vaidya and Johnston, 2013). Notably, decreased HCN conductance correlated with reduced theta resonance in rodents (Marcelin et al., 2009). However, despite the significance of these channels in regulating neuronal excitability and network oscillations, the role of cortical HCN channels in PD is still not fully elucidated.

In this study, we aimed to investigate the potential role of antidromic M2 stimulation during STN-DBS in alleviating motor symptoms in a PD mouse model.

## Results

### Characterization of M2 circuits and behavior in the PD mouse model with medial forebrain lesions with 6-hydroxydopamine (6-OHDA)

The PD mouse model was established by the unilateral stereotaxic administration of 6-OHDA into the medial forebrain bundle (MFB). As expected, we found a substantial reduction in the density of tyrosine hydroxylase (TH)-expressing neurons in the caudoputamen and substantia nigra pars compacta compared with that in the intact hemisphere (Fig. S1A). It was evident that the PD model mice exhibited abnormalities in movement. To attain the behavioral validation of the mouse model, we used a hanging wire test to assess suspension time as mice tried to avoid falling (Darvas et al., 2014; Noor et al., 2016; Shi et al., 2021). The hanging time for control mice increased with repeated tests, likely due to enhanced motor coordination required for hanging. In contrast, 6-OHDA-lesioned mice exhibited a decrease in hanging time, potentially due to the ongoing degeneration of TH neurons (Fig. S1B). We also analyzed the gait pattern of the PD model mice as they walked on a treadmill at a speed of 17 cm/s (Fig. S1C-D). The stance, stride time, and stride length on the contralateral side of the damaged hemisphere were significantly reduced (Fig. S1E). Contrastingly, the stride frequency increased, perhaps to keep up with the speed of the treadmill. The paw area touching the ground varied significantly more than that of control mice, suggesting a less standardized walking pattern.

Having established the PD mouse model, we examined the basal firing rate of the M2 neurons in a PD mouse model by analyzing the single-unit activity (Fig. 1A-B). Consistent with the idea that enhanced M2 neuron activity is involved in the motor-stopping activity in PD symptoms, well-isolated single units (Fig. S2) demonstrated significantly higher firing frequencies with a greater likelihood of burst firing in the M2 of the PD model (Fig. 1D and E).

**Fig. 1.**
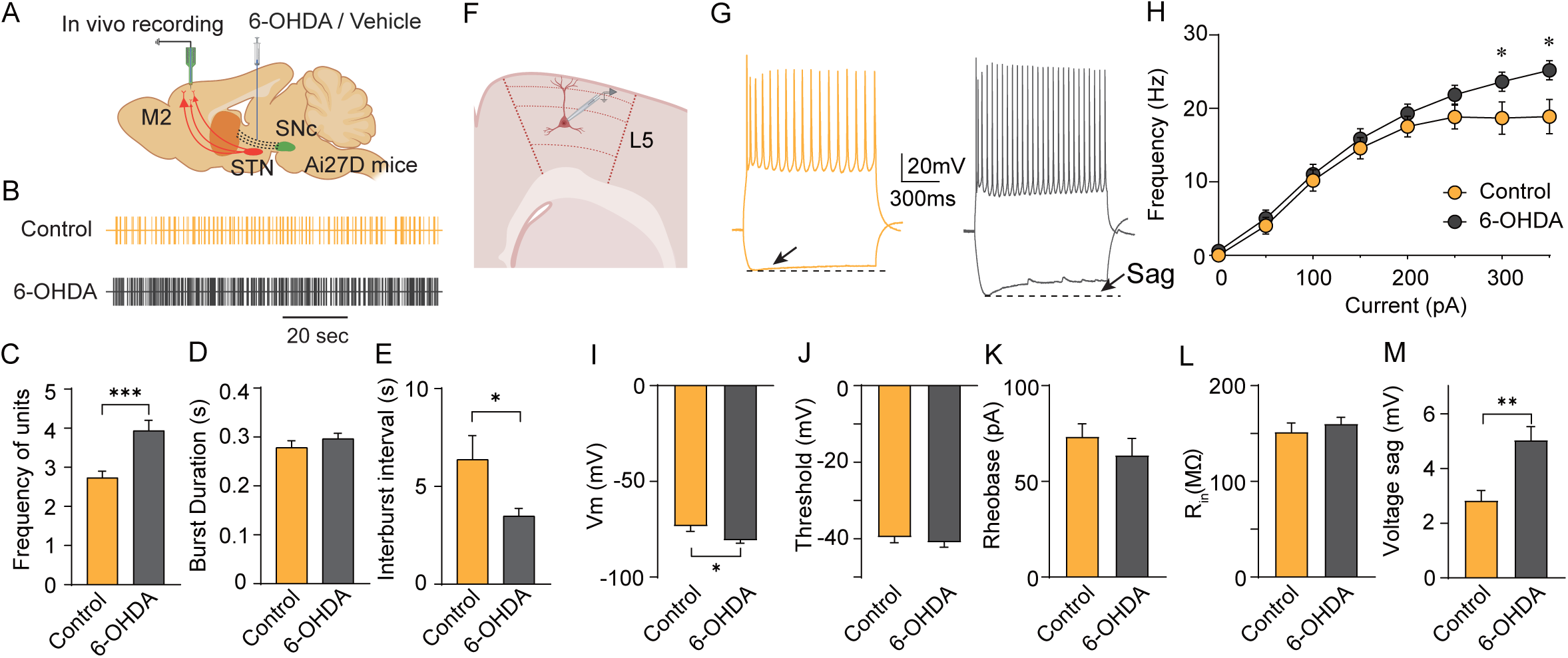
Effect of 6-OHDA lesion on M2 activity. **A.** Schematic illustration of the experiment. To simulate dopamine depletion characteristic of Parkinson’s disease (PD), 6-hydroxydopamine (6-OHDA) was administered into the medial forebrain bundle (MFB) of Ai27D (Rosa-CAG-LSL-hChR2(H134R)-tdTomato-WPR) mice. Basal activities of M2 neurons were assessed via *in vivo* extracellular recordings. **B.** Representative well-isolated units recorded in control and PD mouse models. **C-E.** Population averages of firing rate (left), burst duration (middle), and interburst interval (right) in control (n = 98 neurons) and 6-OHDA-lesioned mice (n = 126 neurons). Spontaneous single-unit activity in the M2 shows higher firing rates in 6-OHDA-lesioned mice compared to controls (firing rate,\*\**p*□=□0.008; burst duration, *p*□=□0.24; interburst interval, \**p*□=□0.044, Mann–Whitney test). **F.** Whole-cell patch-clamp recordings of layer 5 pyramidal neurons in M2 were conducted to assess intrinsic membrane properties. **G.** Representative traces from whole-cell current clamp recordings in M2 layer 5 pyramidal neurons following a 300 pA current injection in control (yellow) and 6-OHDA lesioned (gray) mice. **H.** Averaged action potential frequency as a function of injected current for both control and 6-OHDA lesioned mice. **I.** Resting membrane potential. **J.** Threshold for action potential **K.** Minimal current required to evoke action potential(s), **L.** Input resistance, and **M.** amplitude of voltage sag in control mice and 6-OHDA lesioned mice. Data are presented as mean ± standard error.

Previous studies revealed that fronto-subthalamic inhibitory motor control is mediated by beta-frequency oscillation in humans (Schaum et al., 2021; Sundby et al., 2021), and enhanced synchrony in the beta-frequency range is correlated with bradykinesia in PD patients (Brown, 2007). To test if the enhanced M2 activity is represented in the form of oscillatory synchronous neural activity, we analyzed the local field potential (LFP) in the M2 of the PD mouse model. We found a significant increase in synchronous oscillatory activities over a wide frequency range, with the most dramatic increase in the theta frequency range (Fig. S3).

We then examined whether increased M2 activity can be explained by M2 neuronal membrane property changes in acutely prepared brain slices of PD mouse models (Fig. 1F-M). In corroboration with the increased burst firing and network oscillations (Chan et al., 2004a; Nolan et al., 2007; Robinson and Siegelbaum, 2003), we found the maximum firing frequency by depolarizing current injection was greater in the PD mouse model. Also, the hyperpolarization-activated cation current (I_h_), measured by the amplitude of voltage sag by -200 mV current injection, was significantly enhanced (Fig. 1M). The enhanced excitability in M2 was further established by the enhanced frequency of spontaneously evoked postsynaptic current (sEPSCs) (Fig. S4). These results supported our hypothesis that the enhanced motor-stopping signal, originating from the M2 through a hyperdirect pathway, may contribute to dyskinesia in a PD mouse model.

### Electrical STN stimulation antidromically excites M2 neurons

To test if M2 neuronal activity is involved in the effect of DBS-STN, we first verified the direct synaptic connections between neurons in the M2 and the STN. We delivered retrograde adeno-associated virus (rAAV-retro) to the STN of the transgenic mouse line that expresses channelrhodopsin-2 (ChR2) conjugated with tdTomato in a cyclic recombinase (Cre)-dependent manner (Rosa-CAG-LSL-hChR2[H134R]-tdTomato-WPR or Ai27D) (Fig. 2A-B) (Madisen et al., 2012; Tervo et al., 2016). As previously described (Aron et al., 2003; Kamigaki and Dan, 2017; Li et al., 2020; Magno et al., 2019), we found that a considerable portion of the layer (L) 5 M2 neurons was retrogradely labeled, suggesting direct synaptic connections to the STN (Fig. 2B). Furthermore, we observed a prompt increase in multiunit activity in M2 neurons upon electrical stimulation, imitating DBS-STN (Fig. 2C-D). These results suggest the modulation of M2 neuron activity by DBS-STN and its possible involvement in the effects of DBS-STN.

**Fig. 2.**
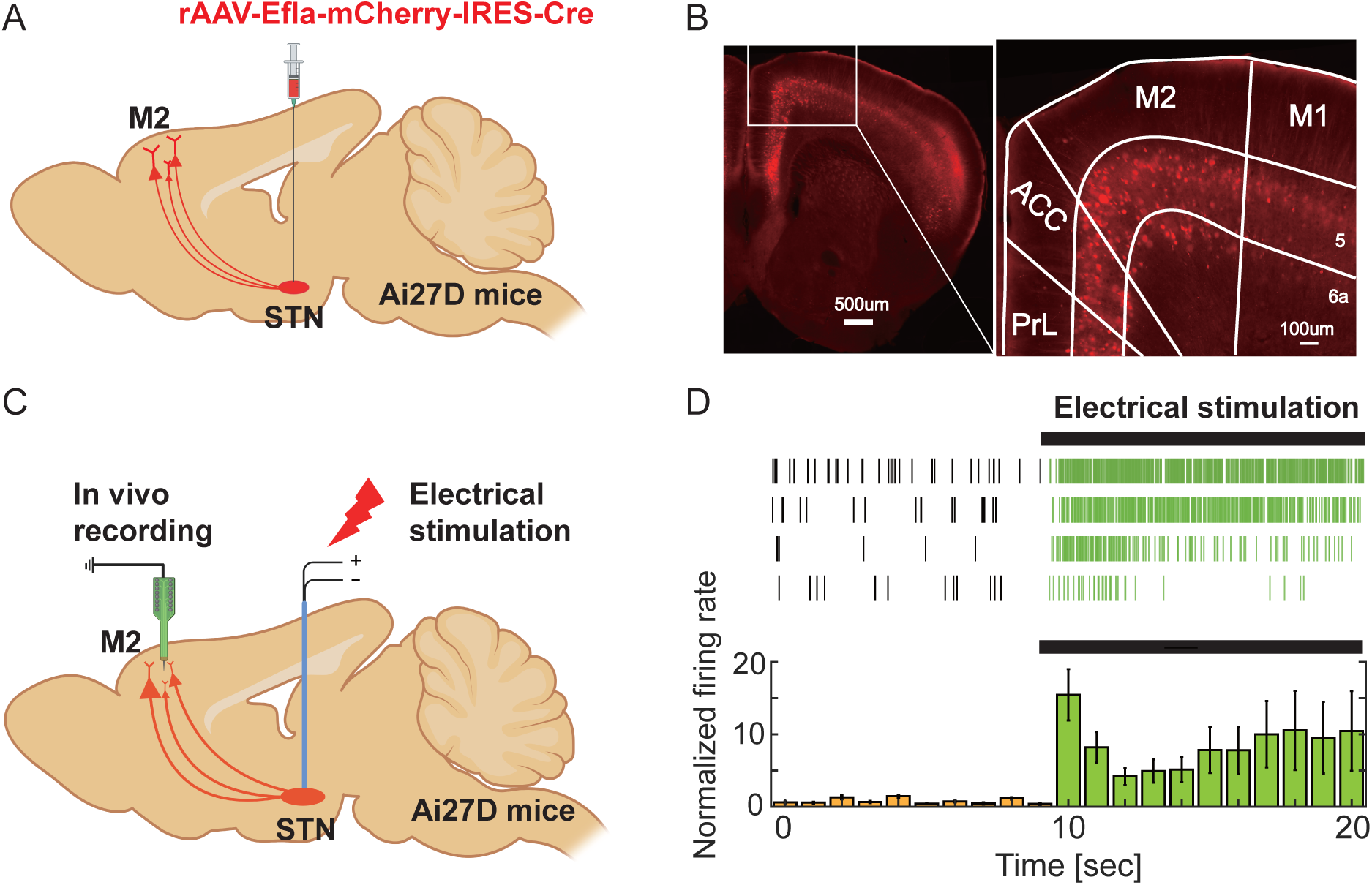
Neurons in the M2 are retrogradely activated by the stimulation of subthalamic nucleus (STN). **A.** Schematic of labeling STN-projecting neurons. rAAV-retro-Ef1a-mCherry-IRES-Cre (≥ 7 x 1012 vig/mL) was injected into the STN of Ai27D mice to trace STN-projecting neurons. **B.** Representative image of STN-projecting neurons in a 50 μm prefrontal brain slice (left, scale bar = 500 μm), and a zoomed-in view of the boxed area (right, scale bar = 200 μm). Labeling includes M1 (primary motor cortex), ACC (anterior cingulate cortex), and PrL (prelimbic cortex). **C.** Schematic diagram illustrating experimental setup for *in vivo* recordings. **D.** Examples (top) and averaged multiunit activity normalized to the mean baseline (bottom) in response to 140 Hz electrical stimulation in the STN.

Prior studies have demonstrated that M2 neurons projecting to the STN are involved in suppressing impulsive behaviors in mice (Kamigaki and Dan, 2017; Li et al., 2020). We explored whether strong activation of deep-layer M2 neurons could also lead to the surprise-induced inhibition of impulsive behavior, akin to the results seen with optogenetic activation of the STN (Fife et al., 2017). In our experiments, an unexpected loud noise was played to water-restricted mice as they licked from a spout. In response, the mice ceased licking in some trials, indicating a potential interruption of their impulsive actions due to surprise. In the trials where the mice ceased licking, we observed a significant increase in the firing rate of M2 neurons (Fig 3). Conversely, there were no changes in M2 activity in trials where licking behavior remained unchanged (Fig. S5), suggesting that M2 neurons indeed positively involved in halting prepotent behaviors, potentially through activating the STN.

**Fig. 3.**
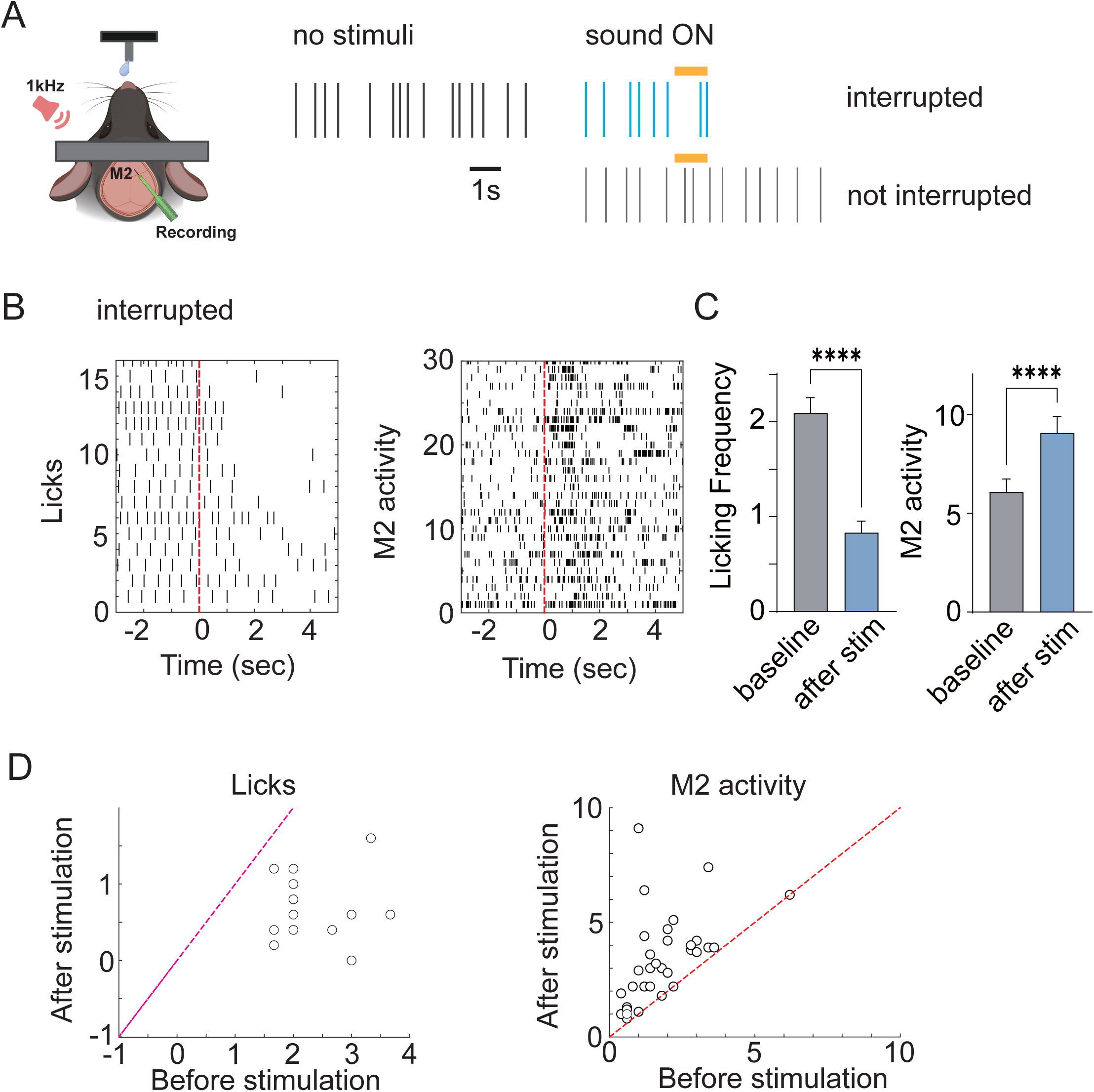
M2 activity upon surprise-induced licking interruptions A. Water-restricted mice were given free access to water, during which licks and M2 activities were recorded. An 85 dB noise was randomly applied for 1 second to assess changes in M2 activity due to surprise-induced licking interruptions. Vertical lines indicate licks. **B.** The raster presentation of licks and simultaneously recorded M2 activity in exemplified trials. **C.** Averaged frequencies of licking and M2 activity of successfully interrupted trials. **D.** Scatter plots of licking an M2 activity frequencies before and after interruption.

### High-frequency stimulation of STN-projecting M2 neurons and the impact on network synchrony and behavioral abnormalities in the PD mouse model

If antidromic stimulation of M2 is involved in the DBS-STN mechanism, the stimulation of STN-projecting M2 neurons should mimic the effect of DBS-STN to a certain extent. To test this, we analyzed animal behavior upon optogenetic stimulation of STN-projecting M2 neurons (Fig. 4). Given the relevance of the theta power to forelimb strength and stride pattern associated with gait freezing and voluntary gripping in PD patients (Shine et al., 2014; Tan et al., 2013), we conducted wire-hanging tests and gait analysis. The observed abnormalities in forelimb strength and stride pattern (Fig. S1) significantly improved immediately after optogenetic stimulation of STN-projecting M2 neurons (Fig. 4, 473 nm, 9 mW/mm^2^, 2 ms square pulses at 140 Hz, 3 min). We found that the latency of falling from the wire significantly increased in response to optogenetic stimulation (Fig. 4B). Likewise, the gait pattern of 6-OHDA-lesioned mice was remarkably normalized (Fig. 4C-E). Specifically, optogenetic stimulation led to increased stance time, stride time, and stride length, as well as a decrease in stride frequency and the variability of the paw area.

**Fig. 4.**
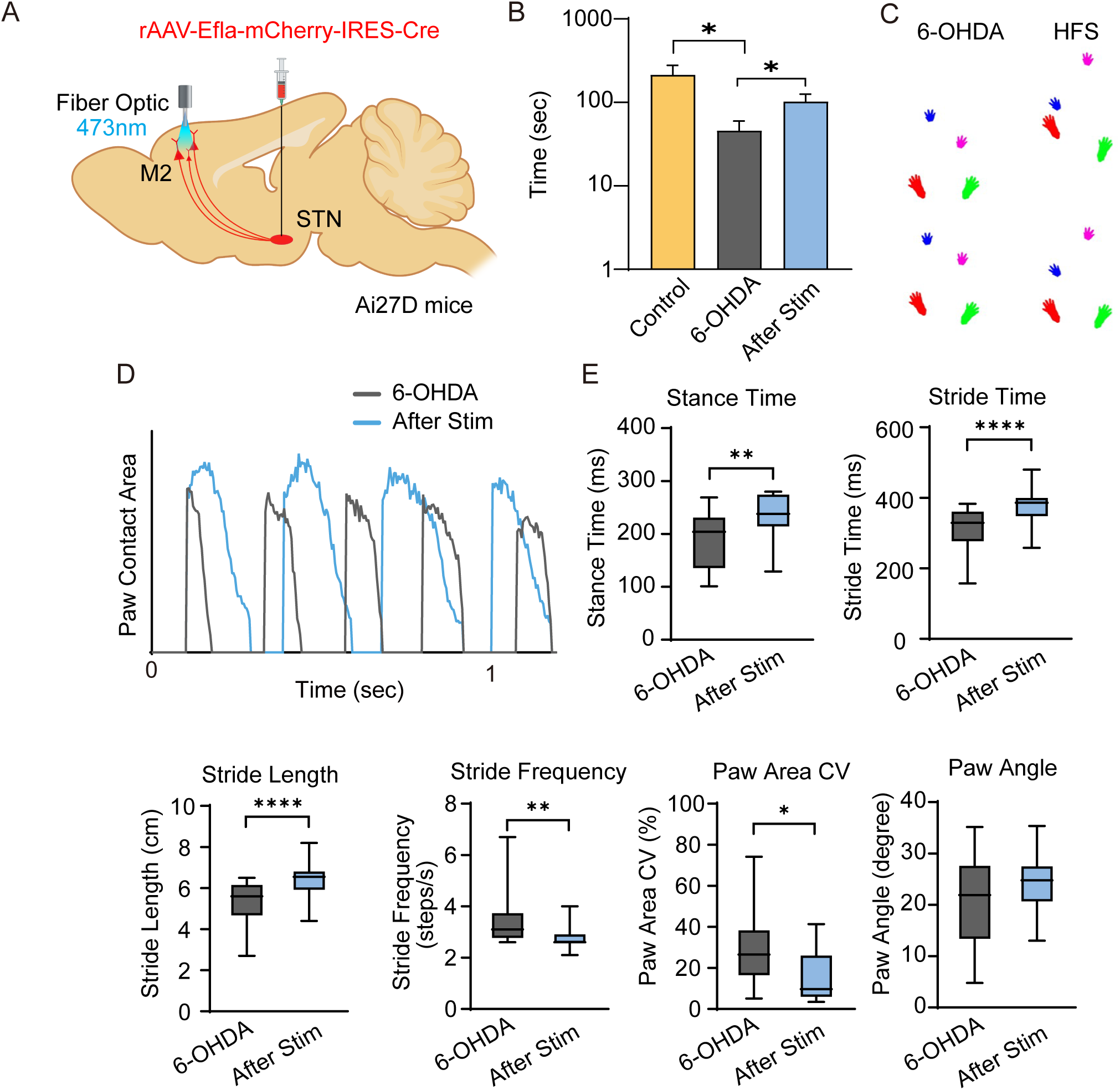
Analysis of motor function in PD mouse models upon optogenetic stimulation of STN-projecting M2 neurons. **A.** Diagram detailing the method for selective stimulation of STN-projecting M2 neuron. **B.** Mean wire-hanging time of PD model mice before and after optogenetic M2 stimulation. Optogenetic stimulation significantly increases the hanging time in the wire hanging test (473 nm, 9 mW/mm^2^, 2 ms square pulses at 140 Hz, n = 7, *p*□=□0.026, paired t-test). **C.** Representative positions of the paws extracted from the ventral plane videography. PD mouse model before (gray) and after (blue) optogenetic stimulation. **D.** Paw area with the left hind limb in contact with the treadmill surface before (dark gray) and after (blue) optogenetic stimulation in 6-OHDA lesioned mice. **E.** Comparison of gait parameters (stance time, stride time, stride length, stride frequency, coefficient of variation (CV) of the paw area, and paw angle) of the paws with the left hind limb of 6-OHDA lesioned mice before and after optogenetic stimulation (****p<0.001, **p<0.01, *p<0.05 relative to 6-OHDA lesioned mice; paired t-test, n =14). Data are presented as box and whisker plots, where each dot represents the mean before and after optogenetic stimulation.

So far, our results indicate that in the PD model, M2 neurons, which are involved in halting prepotent behaviors, exhibited hyperexcitability along with increased oscillatory synchrony. Furthermore, high-frequency stimulation of STN-projecting M2 neurons alleviated behavioral abnormalities in the PD model in a manner similar to DBS-STN (Li et al., 2012b; Schor et al., 2022). We then examined whether the observed improvement in behavioral abnormalities was associated with electrophysiological normalization in M2 neurons. Indeed, optogenetic stimulation of STN-projecting M2 neurons significantly reduced oscillatory synchrony (Fig. S6). Particularly, a reduction in oscillatory synchrony was observed in the theta frequency range (Fig. S6G). Furthermore, optogenetic stimulation modulated the frequency of well-isolated single units recorded in the M2 (Fig. 5). Although frequency modulation varied widely, stimulation reduced the firing frequencies and burst firing duration of the M2 units in the population, with significant effects lasting several minutes after the stimuli ended (Fig. 5). To further examine whether baseline activity may explain the direction of modulation, we grouped units depending on frequency modulation direction (Fig. 5G, black and red circles). The units whose frequency was decreased by optogenetic stimulation had significantly higher baseline activity. In contrast, units with moderate baseline activity showed no changes or a slight increase in activity. Also, the magnitude of firing frequency reduction by the stimulation correlated with pre-stimuli baseline activity (Fig. 5G). The two groups of units had comparable kinetic properties, suggesting that STN-projecting M2 neurons are not distinguishable by cell type (Fig. S7). Thus, our findings suggested that optogenetic stimulation of STN-projecting M2 neurons relieved the hyperactivity of the M2 circuit, and that modulation efficacy may differ depending on baseline activity levels.

**Fig. 5.**
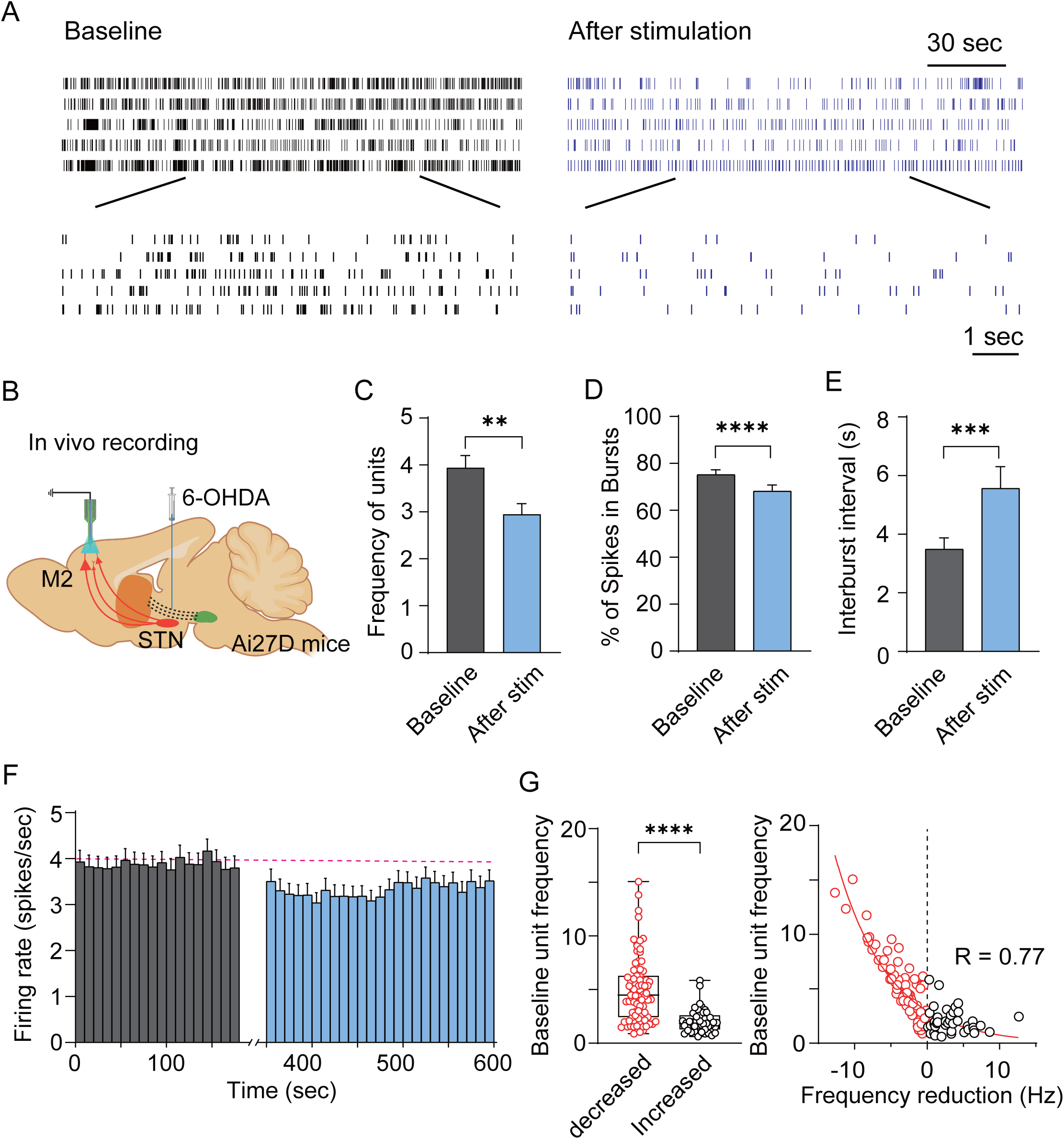
Pattern and frequency of single unit activities measured in the M2. **A.** Example raster plots of representative single units before (left) and after (right) optogenetic stimulation. **B**. Schematic diagram of the experiment. **C.** Changes in spike frequency in M2 upon optogenetic stimulation. **D.** Proportion of spikes occurring in bursts out of total spikes recorded in M2 **E.** Duration between bursts in M2 units following optogenetic stimulation (n= 126 neurons, ****p <0.0001, ^***^p <0.001, ^**^p < 0.01, paired t-test). **F.** Averaged unit counts binned for 10 s from a PD mouse model (n = 126, 19 mice). Baseline counts are color-coded in dark gray and post-stimulation counts are shown in blue **G.** Baseline unit counts grouped by direction of the frequency changes. Baseline activities of units are categorized based on their response to subsequent interventions: inhibited units (n = 83) show significantly higher baseline activity compared to increased units (n = 43). Statistical analysis confirms significant differences (****p < 0.0001, Mann–Whitney test).

### The impact of the high-frequency electrical STN stimulation on M2 neuron activity

The effect of optogenetic stimulation of STN-projecting M2 neurons may differ from antidromic M2 activation during electrical STN stimulation, considering factors such as fidelity of antidromic stimulation, difference in action potential kinetics, and synaptic transmission (Chomiak and Hu, 2007; Jackman et al., 2014). To directly address this potential difference, we examined whether the amelioration of M2 hyperactivity may be replicated by electrical STN stimulation (Fig. 6). The stimulation electrode was placed in the electrophysiologically identified STN, while the activity of M2 neurons was monitored extracellularly before, and after stimulation. In line with the optogenetic stimulation findings, isolated M2 unit frequencies were significantly reduced by electrical stimulation (Fig. 6C, S8). Contrastingly, the M2 neuron frequencies showed the opposite effect in the healthy hemisphere (Fig. S8) or in control mice (Fig. 6D), suggesting that the M2 frequency reduction is specific to PD-like symptoms.

**Fig. 6.**
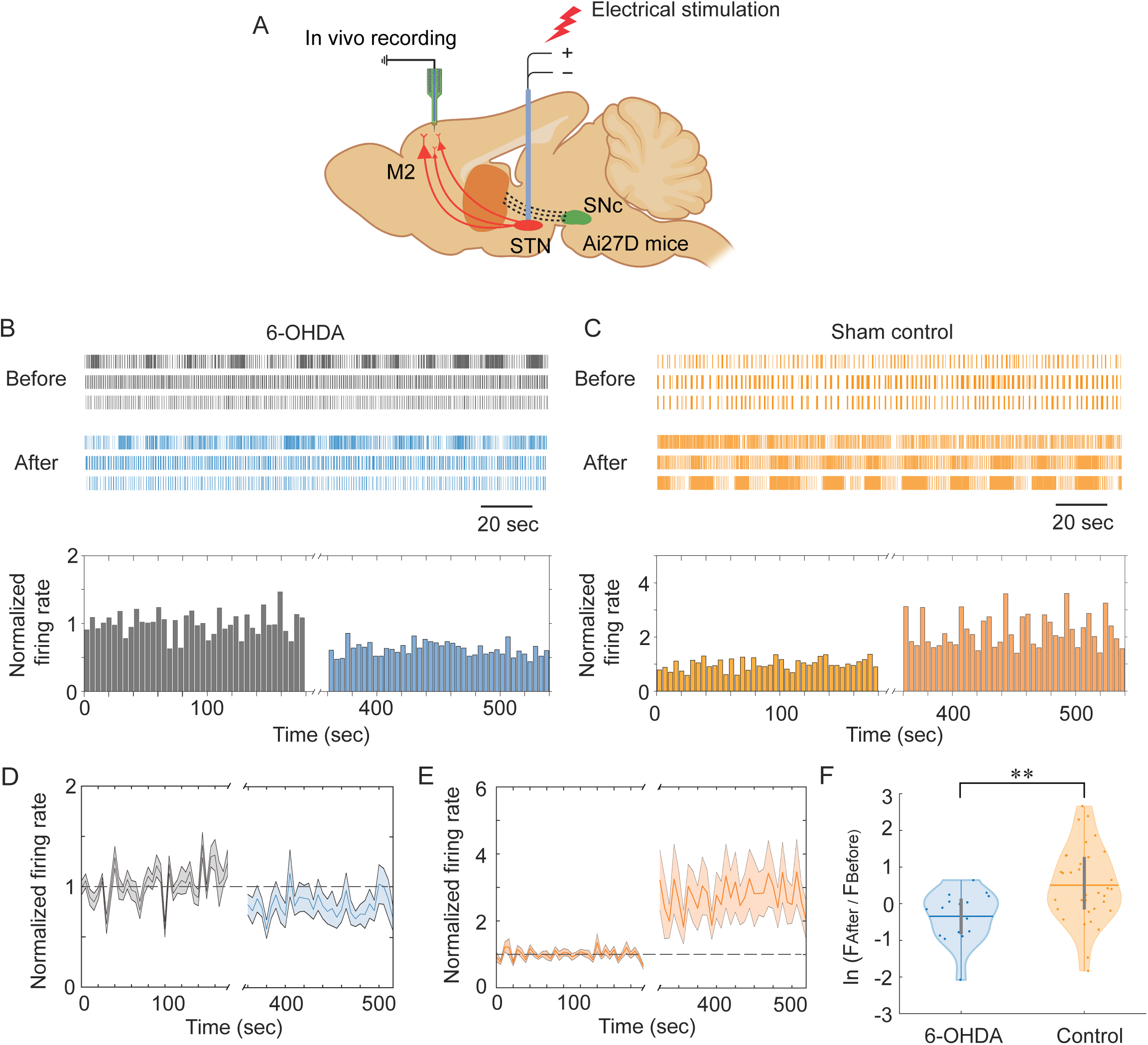
Frequency of single units recorded in the M2 in response to electrical STN stimulation. **A.** Schematic diagram of the experiment. **B-C.** Raster plots (upper) and its 4.5-s time-binned histogram (lower) of three representative single units in the M2 for 6-OHDA (B, n = 16) and sham control (C, n = 35) mice. The firing rates are normalized to mean firing of the baseline activities. **D-E.** Moving averaged (5-s time window without overlapping) firing rate of all single units measured from 16 OHDA-lesioned mice (D) and 35 sham control mice (E). The firing rate is normalized to the mean firing rate before the stimulation. In both plots, the dashed gray line illustrates the normalized firing rates before stimulation. The solid lines represent the average firing rates. The shaded area represents the standard error of the mean for all units for each condition. **F**. Violin plots of firing rate ratio before and after stimulation. A single scatter represents the firing rate of a single unit using a natural logarithm. The thick vertical lines show the first and third quartiles, whereas the thin vertical lines display 95% confidence intervals. The white dots and thick horizontal lines represent the median and mean values, respectively (n = 16, PD model and n = 35, control; ** p<0.01, unpaired *t*-test).

### M2 neural excitability upon optogenetic M2 stimulation

To explore the cause of the reduced M2 neurons firing, we analyzed the membrane properties of M2 neurons of 6-OHDA-lesioned mice upon optogenetic stimulation *ex vivo*. Approximately four weeks after the MFB lesion, when the behavioral symptoms became noticeable, we measured M2 neuronal excitability before and after optogenetic stimulation with whole-cell recordings. Corroborating our *in vivo* observations, we found a significant reduction in excitability by optogenetic stimulation of STN-projecting M2 neurons. Significantly fewer action potentials were evoked in response to depolarizing current injections at all amplitudes examined (Fig. 6A-C). Despite a slightly depolarized resting membrane potential (Fig. 6C) and a comparable threshold (Fig. 6D), a significantly larger depolarizing current was required to evoke an action potential (Rheobase, Fig. 6E). We attributed this reduced excitability to the reduced input resistance (Fig. 6F) and hyperpolarization-evoked depolarizing voltage sag (Fig. 6G).

Previously, it has been shown that, during DBS-STN stimulation, antidromically stimulated M1 neurons temporally suppressed M1 circuits through activation of inhibitory neuron (Valverde et al., 2020). To determine if a similar pattern of circuit suppression occurs in M2, we examined spontaneously evoked excitatory and inhibitory synaptic currents (sEPSC and sIPSC, respectively). If similar circuit suppression was involved, enhanced sIPSC and reduced sEPSC frequency would be expected. However, we did not find any significant alterations in spontaneous synaptic frequencies (Fig. S9), suggesting that the reduced excitability of M2 neurons is unlikely to be mediated by the same circuit mechanism as M1.

### HCN channel dependency of high-frequency M2 stimulation-induced behavioral amelioration

Considering the I_h_ (Fig. 1M), increased firing frequency, burst firing (Fig. 1 C and E) and theta-band oscillation (Fig. S3) observed in the PD model, along with their reversal through STN stimulation (Figs S3, 5, 7, S3 and S6), we ought to test whether concurrent changes in the I_h_ current could account for these alterations in neural activity and behavioral improvement (Chan et al., 2004b; Nolan et al., 2007; Robinson and Siegelbaum, 2003). To examine the behavioral impact of reduced HCN channel activity in M2, we administered ZD7288, a selective HCN channel blocker, into the M2 region of PD mice, and subsequently assessed the effects on behavioral abnormalities (Fig. 8A). Remarkably, we observed a significant improvement in behavioral abnormalities, as evidenced by enhancements in wire-hanging time and gait patterns (Fig. 8). These findings strongly suggest that the observed behavioral improvement upon M2 neuron stimulation can be attributed to reduced HCN channel activity.

## Discussion

In this study, we explored the possibility that repetitive antidromic activation of M2 neurons during DBS-STN may be involved in the effects of DBS. We hypothesized that hyperexcitability and normalization of the STN and its upstream premotor areas, which are associated with motion-stopping (Adam et al., 2022; Aron et al., 2007b; Eagle et al., 2008; Emmi et al., 2020; Li et al., 2020; Mi et al., 2019; Schaum et al., 2021; Sundby et al., 2021; Swann et al., 2012), contribute to the motor dysfunction observed in PD, while the normalization of these areas through DBS-STN alleviates motor symptoms. In line with previous reports (Adam et al., 2022; Kamigaki and Dan, 2017; Li et al., 2020; Magno et al., 2019), we found that a notable population of L5 neurons in M2 project directly to the STN (Fig. 2) and M2 neurons are activated by unexpected loud noise (Fig. 3). This demonstrates the potential involvement of M2 neurons during DBS-STN. To isolate the effect of high-frequency M2 activity during DBS-STN, we used optogenetic stimulation of STN-projecting M2 neurons (Fig. 4A). High-frequency stimulation of these neurons, simulating DBS, alleviated the observed abnormalities in behavior (Fig. 4) and oscillatory synchrony (Fig. S6) in the PD mouse model. The causality of network oscillations with motor impairment was not directly addressed in this study. However, oscillatory activity has been observed to be repeatedly correlated with the level of motor impairment in both patients with PD and animal models (Dorval et al., 2010; Levy et al., 2002). Furthermore, the premotor areas directly connected to the STN have been indicated as the source of oscillation during stop-signal responses (Schaum et al., 2021; Wagner et al., 2018). Thus, we hypothesized that the hyperexcitability of STN-projecting M2 neurons could be a source of dyskinesia in PD.

Consistent with our hypothesis and observed hyperactivity in M2 of PD patients (Buhmann et al., 2004; Samuel et al., 1997), we found that the STN-projecting M2 neurons fired at significantly higher frequencies in the PD model than in the sham-control mice (Fig. 1), which can be suppressed upon optogenetic stimulation accompanied by behavioral amelioration (Fig. 4 and 5). The reduced activity of M2 neurons upon optogenetic stimulation could be recapitulated *ex vivo*, where we found a significant reduction in the firing frequency in response to the depolarizing current injection (Fig. 7). This phenomenon could be attributed to the reduced R_in_, which requires greater excitatory inputs to fire an action potential. Moreover, diminished I_h_ current, the hallmark of the HCN channel, could contribute to reducing the repetitive firing of neurons (Fig. 7) (Chan et al., 2004b; Thuault et al., 2013; Yang et al., 2018). Consequently, our results demonstrated that reducing HCN channel activity in M2 with ZD7288 mimicked the effects of DBS-STN to some extent (Fig. 8), suggesting that HCN channels could serve as potential therapeutic targets for PD.

**Fig. 7.**
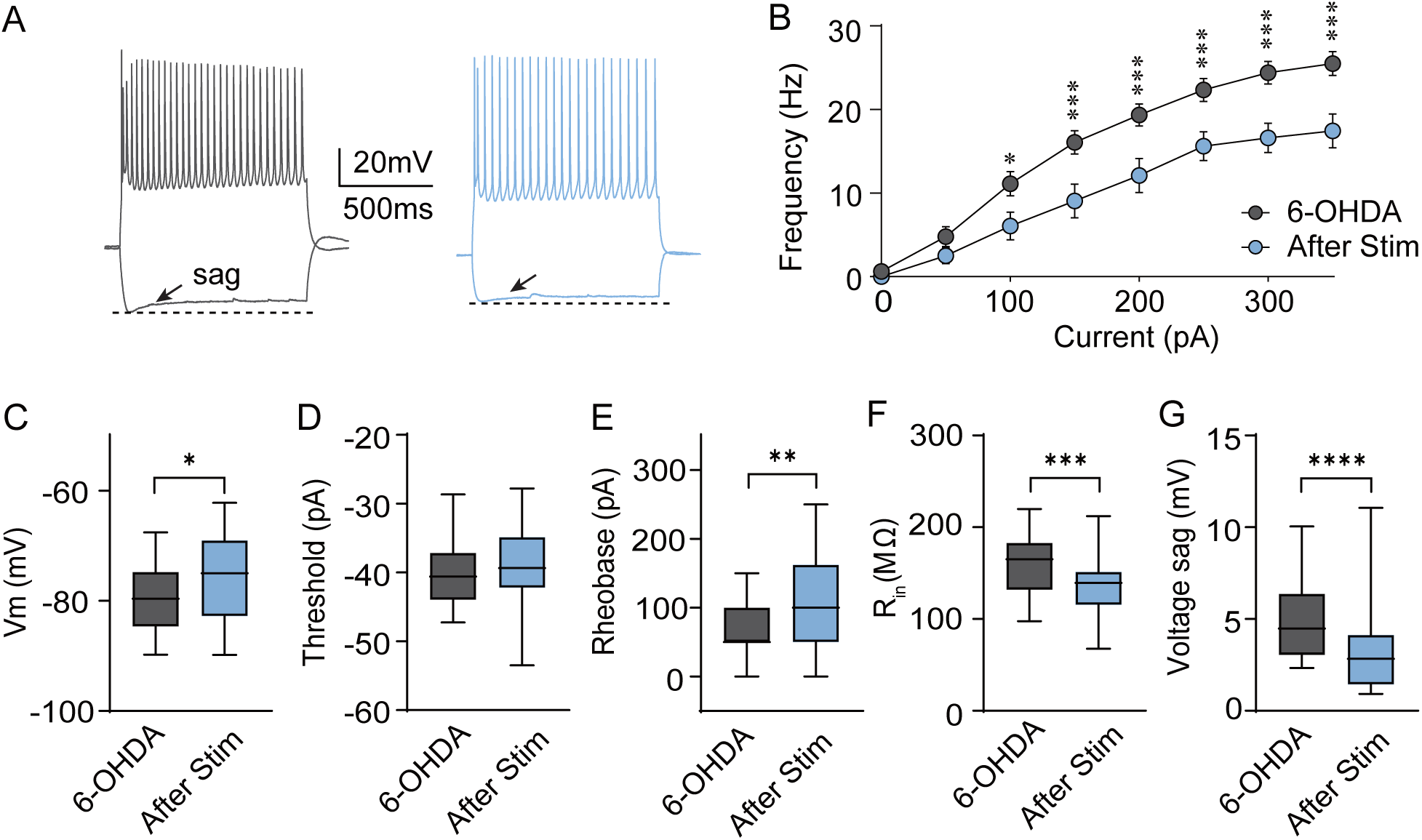
Intrinsic excitability of M2 neurons upon optogenetic stimulation in the PD mouse model. **A.** Example voltage traces of M2 neurons in 6-OHDA mice in response to depolarizing current injections at -200 and 350 pA amplitudes. **B.** Mean firing frequencies of 50-350 pA depolarizing step current injections before (dark gray) and after (blue) optogenetic stimulation. **C.** Resting membrane potentials (-79.49 ± 1.44 mV before stimulation vs. -75.24 ± 1.92 mV after stimulation, n = 18, p = 0.014, paired t-test). **D.** Action potential threshold (-40.36 ± 1.14 mV before and -39.35 ± 1.38 mV after stimulation, n = 18, p = 0.19, paired t-test). **E.** The minimum current required for evoking an action potential, rheobase (66.66 ± 9.90 pA before and 113.88 ± 16.57 pA after stimulation, n = 18, p = 0.002, paired t-test), **F.** Input resistance (before stimulation, 159.12 ± 8.13 mV vs. 134.15 ± 7.52 mV after optogenetic stimulation, n = 18, p = 0.0005, paired t-test), and **G.** voltage sag (4.90 ± 0.53 mV before and 3.46 ± 0.62 mV after stimulation, n = 18, p = 0.0018, paired t-test). Data are presented as box and whisker plots, where each graph shows the mean before and after optogenetic stimulation.

**Fig. 8.**
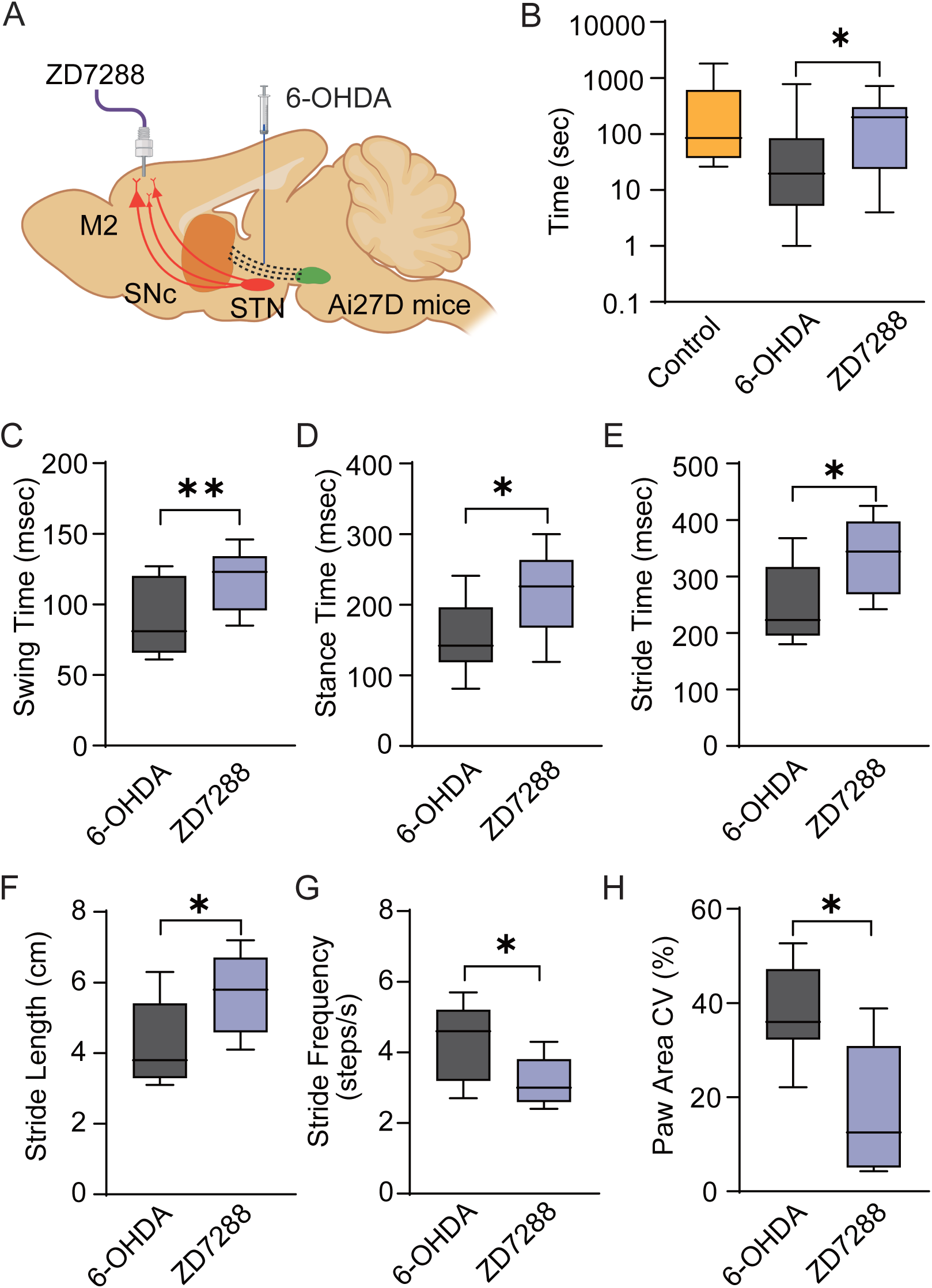
Effects of ZD7288 microinjection into M2 on the motor abnormalities in the PD mouse model. **A.** Schematic illustration of the experiment. **B.** Mean wire-hanging time of control, PD model mice before and after ZD7288 administration. Wire-hanging time is increased after microinjection of ZD7288 in the PD mouse model (n = 12, *p*□=□0.031, paired t-test). **C-H**. Comparison of gait parameters (swing time, stance time, stride time, stride length, stride frequency, coefficient of variation (CV) of the paw area) of the paws with the left hind limb of 6-OHDA lesioned mice before and after ZD7288 administration (**p<0.01, *p<0.05 relative to 6-OHDA lesioned mice; paired t-test, n =7). Data are presented as box and whisker plots, where each box represents the mean before and after ZD7288 administration.

### Oscillatory network activity in the M2

Typically, enhanced beta-frequency oscillations are reported to correlate with movement abnormalities in the STN of patients with PD (Brown, 2007). However, in studies using various animal models, the increase in beta frequency was not consistent. Instead, synchronous oscillations in lower-frequency ranges, such as theta, were observed (Devergnas et al., 2014; Leblois et al., 2007; Niyomrat et al., 2021; Soares et al., 2004; Stein and Bar-Gad, 2013; Wang et al., 2017). Similarly, enhanced oscillatory power in the lower-frequency range, including the theta band, was observed in some studies of patients with PD (Moazami-Goudarzi et al., 2008; Morita et al., 2009; Shine et al., 2014; Stoffers et al., 2007; Tan et al., 2013). Increased synchrony in the theta range in the STN and pre-SMA was correlated with gait freezing and hand use (Breit et al., 2007; Hutchison et al., 1997; Shine et al., 2014; Tan et al., 2013). Thus, it is feasible that enhanced synchronous neural activity itself is the pathological hallmark of PD, whereas the abnormal frequency range is related to the type of motor symptoms (Stein and Bar-Gad, 2013; Tan et al., 2013). Correspondingly, we observed gait abnormalities and shortening of the hanging time, along with the power of oscillation in all frequency ranges, the greatest being in the theta range (Fig.S1 and S3). Furthermore, we found that selective stimulation of STN-projecting M2 neurons significantly reduced the theta frequency range, together with the behavioral symptoms, of PD mice (Fig. 4 and S6). The observed enhancement of oscillatory synchronization in the M2 region of the PD mouse model, as well as its normalization through M2 stimulation, corresponded to concurrent changes in I_h_ current (Fig. 1 and 7). Distally enriched distribution and inductance by HCN channels synchronize the distal and somatic voltage waveform, with maximum synchronization occurring within the theta frequency range (Hu et al., 2009; Narayanan and Johnston, 2008; Pike et al., 2000; Rotstein et al., 2005). While previous studies have primarily investigated hippocampal pyramidal neurons, similarly polarized HCN distribution in neocortical pyramidal neurons suggests a potential for a similar synchronization effect (Kim et al., 2022; Paxinos and Franklin, 2001).

### Antidromic STN-projecting neurons during DBS-STN

The activity of STN neurons is dependent on many brain areas in addition to the M2. Notably, M1 involvement has attracted considerable attention. High-frequency optogenetic stimulation of M1 neurons, or afferent axons from the M1, improved the behavior efficiently (Gradinaru et al., 2009; Sanders and Jaeger, 2016). A recently published study showed that DBS-STN relieves the M1 circuits from hyperexcitation by activating SST-expressing inhibitory neurons in the M1 (Valverde et al., 2020). We tested whether a parallel phenomenon occurs in the M2 by measuring the frequencies of sEPSCs and sIPSCs. We could not detect any significant frequency changes after high-frequency stimulation of the STN-projecting M2 neurons (Fig. S9). However, we cannot rule out local inhibitory neuronal involvement in normalized M2 activity if the effect of the inhibitory neuron lasts only during stimulation. In fact, the enhanced activity of SST neurons in the M1 lasted less than a minute (Valverde et al., 2020), which may not be long enough to reliably detect frequency changes in sIPSCs. It is unlikely that the effect of GABAergic synaptic transmission activated by STN-projecting neurons lasts longer than that time scale. It is noteworthy that all behavioral and electrophysiological changes illustrated in our study were measured after stimulation of STN-projecting M2 neurons. Furthermore, feedforward inhibition-mediated circuit silencing is difficult to reconcile with the selectivity effect observed in the M2. We observed that optogenetic stimulation occurred selectively in the set of units with high baseline activity, but not in neurons with low baseline activity or in control mice (Fig. 4 F-G and Fig. 5). Together, we propose that excitability reduction, rather than inhibitory neuron-mediated suppression, is the leading cause of the long-lasting effects of DBS-STN through M2. The observed effect of M2 stimulation in this study was not mutually exclusive from previous studies that implicated the involvement of M1 in the therapeutic effect of DBS-STN. In fact, we believe that the therapeutic effect of DBS-STN could be due to a combination of various circuit modulations, including the M1 and M2, each of which may relate to different effect times and behavioral symptoms.

### Study novelty

Similar to our study, Magno et al. (Magno et al., 2019) found that high-frequency M2 stimulation alleviates behavioral PD-like symptoms in 6-OHDA mice. However, they accounted for this phenomenon via increased dopamine concentration in the striatum. We did not directly measure dopamine replenishment in our study. However, we found no changes in the aperiodic components of the subthalamic LFP (Fig. S10), which have been reported to vary with striatal dopamine concentration (Dastin-van Rijn et al., 2021; Kim et al., 2022). This result suggests that dopamine may not be the primary explanation for the optogenetic M2 stimulation under our experimental conditions. This apparent discrepancy can be explained by differences in the population of stimulated M2 neurons. Magno et al. transduced ChR2 into excitatory neurons in all layers of the M2 and stimulated M2 or M2 axons passing the dorsomedial striatum (Magno et al., 2019). Thus, optogenetic stimulation could have influenced wide brain areas, including the ventral tegmental area, the major source of dopaminergic neurons. In this study, we selectively stimulated STN-projecting M2 neurons to study M2 circuit modulation. Therefore, this study directly addressed the therapeutic contribution of antidromic M2 stimulation to DBS-STN.

### Limitations of the study

While M2 activity modulation has therapeutic potential, several essential issues remain. First, future research should determine whether the reduced activity or decreased excitability is sufficient to ameliorate PD-like symptoms of the mouse model. Furthermore, it remained unclear how dopamine depletion causes hyperexcitability in the M2 region of the PD mouse model. The optimal activation frequency of the STN-projecting M2 neurons needed to achieve the therapeutic effects remained unclear in this study. Although we observed improved behavioral and electrophysiological symptoms using 140-Hz optogenetic stimulation, the actual firing frequency and duration of the STN-projecting M2 neurons could not be determined because of the low fidelity action potentials by high-frequency ChR2 stimulation. Likewise, while we observed a reduced firing rate of the M2 neurons by antidromic electrical stimulation with the same frequency at the STN, somatic firing frequency should be significantly lowered by voltage filtering through axons (Chomiak and Hu, 2007; Jackman et al., 2014). Thus, a systematic comparison with different stimulation frequencies may be required.

### Significance and future perspectives

Our study has important implications and opens new avenues for future research. First, our findings provide a scientific basis for the development of less invasive approaches for stimulating the frontal cortices that project to the STN, while still preserving the therapeutic effects DBS-STN (Mi et al., 2020, 2019; Sundby et al., 2021). This could potentially lead to the development of novel stimulation techniques that are more targeted and have fewer side effects. Furthermore, our study highlights the potential of targeting excessive activity in the M2 region as a therapeutic strategy for alleviating motor symptoms in PD. The significant improvement in motor abnormalities observed with ZD7288 in our PD mouse model suggests that these channels could serve as promising drug targets for PD treatment. Further exploration of HCN channel blockers and their effects on PD symptoms in clinical settings is warranted.

Overall, our findings shed light on the mechanisms underlying the therapeutic effects of DBS-STN and present new opportunities for the development of less invasive stimulation techniques and potential drug therapies for PD. Further investigations into these areas hold great promise for improving the quality of life for individuals living with PD and advancing our understanding of the neural circuits involved in motor control.

## Materials and Methods

### Animals

Transgenic mice expressing the ChR2 variant, ChR2 H134R, fused to tdTomato were generated by crossing heterozygous B6.Cg-Gt(ROSA)^26Sortm^ ^27.1(CAG-COP4*H134R/tdTomato)Hze^/J (Ai27D; Jackson Laboratory, Bar Harbor, ME, USA; stock number 012567) and C57BL/6J (Jackson Laboratory, Bar Harbor, ME, USA; stock number 000664) mice. Mice were placed in a 12 h reversed light/dark cycle (lights on at 8 p.m.) with ad libitum access to standard chow and water. All animal experiments were approved by the Institutional Animal Care and Use Committee of the Korea Brain Research Institute (IACUC 22-00009).

### Stereotactic surgery

A total of 81 male mice were randomly assigned to the following two groups: 6-OHDA lesioned mice (n = 66) and control mice (n = 15). Mice were anesthetized with a ketamine-xylazine mixture of 100 mg/kg ketamine-10 mg/kg xylazine and positioned in a stereotaxic apparatus (RWD Life Science Inc., San Diego, CA, USA). Body temperature was controlled throughout the procedure using a feedback heating pad. Thirty minutes prior to intrastriatal injection, mice received desipramine (10 mg/kg intraperitoneal injection [i.p.], Sigma, St. Louis, MO, USA) to protect noradrenergic terminals against 6-OHDA-induced damage.

We injected 6-OHDA hydrochloride (Sigma, St. Louis, MO, USA) to unilaterally lesioned dopamine neurons (using a 10 μL NanoFil syringe [WPI, Sarasota, FL, USA] connected to an automated pump at 80 nL/min [UltraMicroPump and SYS-Micro4 Controller, WPI, Sarasota, FL, USA]) in the MFB [anteroposterior (AP): -1.1 mm; mediolateral (ML): -1.1 mm; dorsoventral (DV): -4.7 mm; coordinates from Paxinos and Franklin, 2001 (Paxinos and Franklin, 2001) of middle-aged adult mice (10-12 weeks). The 6-OHDA animals were injected with 300 nL of 6-OHDA hydrochloride (5 μg/μL) with 0.2 mg/mL ascorbic acid in 0.9% NaCl, while the control animals received the same volume of vehicle solution at the same coordinates. This method produced strong nigrostriatal lesioning, as verified histologically through TH staining.

In the same surgery, adeno-associated viruses carrying a retrograde tracer, Cre construct (Efla-mCherryIRES-WGA-Cre), were injected in the STN (AP: -1.8 mm; ML: -1.56 mm; DV: -4.3 mm from the bregma). Chronic optical fiber implants were manufactured in ceramic ferrules with 200µm inner diameter (RWD Life Science Inc., San Diego, CA, USA) in the M2 (AP: 1.7 mm; ML: -0.6 mm). Ketoprofen (5 mg/kg) was injected subcutaneously once daily for three days. Mice were allowed to recover for 4 weeks after virus injection to allow axonal expression of opsin.

For experiments, HCN channel blocker microinjection, a stainless-steel guide cannula (26-gauge stainless steel wire, Plastics One, Roanoke, VA) was implanted into the M2 (AP:1.7 mm; ML: -0.6 mm), and a dummy cannula (33-gauge stainless steel wire, Plastics One, Roanoke, VA) was inserted into the guide cannula to reduce the incidence of occlusion. The guide cannula was fixed to the skull using a super-bond. HCN channel blocker, ZD7288 (Tocris, Bristol, UK) solutions was infused into the M2 using a microinjection unit (33-gauge cannula) that attached to a 10 μL NanoFil syringe via polyethylene tubing (PE-50). ZD7288 was dissolved in sterile saline and infused into the M2 (300nl, 80 nl / min).

### Electrophysiology

#### In-vivo recording

For *in-vivo* electrophysiological experiments, mice were anesthetized with ketamine (100 mg/kg, i.p.) and placed in a stereotaxic device (WPI, Sarasota, FL, USA). In all surgical preparations, the scalp was incised and holes were drilled in the skull overlying the site of recording or stimulation, according to the coordinates from the atlas of Paxinos and Franklin (Paxinos and Franklin, 2001): M2: AP: +1.7 mm from the bregma, ML: ± 0.6 mm from the midline, and DV: -0.5-0.7 mm below the dura; STN: AP: -4.3 mm from the bregma, ML: 1.56 mm from the midline at an anteroposterior angle of 30°, and DV: -4.8-5.0 mm below the dura. An additional hole was drilled over the other cortex in the left hemisphere to implant a microscrew that was used as a recording reference. During the recording, the brain surface was kept wet by dropwise addition of saline.

To record M2 responses, 16-channel silicon electrodes with a fiber optic ferrule (200 µm diameter, A1×16-5mm-100-177-OA16LP; NeuroNexus, Ann Arbor, MI, USA) were used. The optoelectrode was gradually lowered until the tip was positioned at a depth of ∼700 Dm under the surface. Light stimulation was provided by a 473 nm laser with a laser power set to 9 mW, as measured at the tip of the fiber (0.22 NA; Doric Lenses, Quebec, Canada), to be connected to the ferrule on the mouse’s head. Neuronal activity was amplified and digitized using a PZ5 NeuroDigitizer and RZ5D BioAmp Processor (TDT, Alachua, FL, USA) with a 24,414 Hz/channel sampling rate and bandpass filtered between 0.3 and 5 kHz. Single units were identified and sorted into clusters by Offline sorter (Offline sorterTM, Plexon Inc., Dallas, TX, USA)) and further analyzed using NeuroExplorer (NEX Technology, Colorado Springs, CO, USA). Viral expression was allowed for a minimum of 4 weeks before behavioral testing.

#### Four-limb hanging-wire test

The four-limb hanging-wire test was performed to examine motor function impairments. Each mouse was placed on the wire lid of a conventional housing cage, which was then turned upside down. The time spent hanging was recorded from the time of inversion until all paws were released from the grid.

#### Automated treadmill gait test

Gait analysis was performed using DigiGait software and a treadmill (Mouse Specifics, Inc., Framingham, MA, USA) from 5 weeks after administration of 6-OHDA. Mice were placed on a motorized treadmill within a plexiglass compartment (∼25 cm long and ∼5 cm wide). Digital video recordings were acquired using a camera mounted underneath the treadmill to visualize paw contacts on the treadmill belt. The treadmill was set at a fixed speed of 17 cm/s, at which most animals were able to move continuously. Mice were allowed to accommodate the movement of the treadmill for several seconds before the gait analysis. The gaits during the first 5 s after the acclimation period were analyzed using DigiGait software, which automatically identifies paw footprints and processes them to calculate values for multiple gait parameters, including stride frequency and length, paw area CV, and paw angle (Fig. S1).

#### The STOP response of unexpected loud sound

Mice were anesthetized with ketamine (100 mg/kg, i.p.). The head-bar (stainless steel) was attached to the head using a super bond. Mice were allowed to recover after head-bar implantation and were water-restricted in their home cage. For head-fixed mice, the water reward was delivered through a water port located within reach of the mouse’s mouth. Mice were habituated by daily handling for 3-4 days prior to task. The licking response was triggered by a custom-made Arduino controlled by a MATLAB Psychtoolbox script. The unexpected loud sound was delivered by noise (85dB, 1s). Licking frequency and M2 activity were measured using a PZ5 NeuroDigitizer and RZ5D BioAmp Processor (TDT, Alachua, FL, USA) and further analyzed using NeuroExplorer (NEX Technology, Colorado Springs, CO, USA).

### Euthanasia

Mice were deeply anesthetized with ketamine (100 mg/kg, i.p.) and transcardially perfused with ice-cold dissection solution or 4% paraformaldehyde in 0.01□M phosphate-buffered saline (PBS). The brains were removed for histological or electrophysiological studies.

### *Ex-vivo* whole-cell recording

To prepare *ex-vivo* slices for whole-cell recording, brains were removed and immersed in an ice-cold cutting solution with the following composition (in mM): choline chloride, 110; KCl, 2.5; NaHCO_3_, 25; NaH_2_PO_4_, 1.25; glucose, 25; CaCl_2_, 0.5; MgCl_2_·6H_2_O, 7; sodium ascorbic acid, 11.6; and pyruvic acid, 3. The measured osmotic concentration was 320–330 mOsm.

We prepared 300 µm thick acute brain slices containing M2 coronally using a VT1200S Vibratome (Leica Biosystems, Deer Park, IL, USA). The composition of artificial cerebrospinal fluid (aCSF) was (in mM): NaCl, 119; KCl, 2.5; NaHCO_3_, 26; NaH_2_PO_4_, 1.25; glucose, 20; CaCl_2_, 2; MgSO_4_, 1; ascorbic acid, 0.4; and pyruvic acid, 2. The osmotic concentration was 305–310 mOsm.

During sectioning, the solutions were oxygenated using 95% O_2_ and 5% CO_2_. After 30 min of recovery time at 32 °C, aCSF slices were transferred to room temperature. Recordings were performed using patch pipettes (3.5–5 MΩ) filled with an internal solution consisting of the following (in mM): KCl, 20; potassium gluconate, 125; HEPES, 10; NaCl, 4; EGTA, 0.5; ATP, 4; TrisGTP, 0.3; and phosphocreatine, 10 (pH 7.2, 290–300 mOsm).

#### Immunohistochemistry

Brains were removed and post-fixed for 12 h at 4 °C. Coronal sections (50 µm) were obtained using a vibratome (VT1000S; Leica, Nussloch, Germany). Slices were immersed in 0.2% Triton X-100 Tris-buffered saline solution (TBS-T) for 10 min, followed by blocking for 1□h with 0.2% Triton X-100 and 5% normal goat serum (Jackson ImmunoResearch, Suffolk, UK) in PBS. Sections were washed in TBS-T and incubated overnight at 4 °C in the same buffer with anti-TH monoclonal antibodies (1:500; Millipore, Stockholm, Sweden). Immunoreactivity was detected using a secondary antibody conjugated with Alexa fluorescent dye (1:400; Jackson Laboratory, Bar Harbor, ME, USA) for 3 h at room temperature. The slices were mounted on glass slides with a mounting solution containing DAPI (4’,6-diamidino-2-phenylindole; Vectashield with DAPI, Vector Laboratories, Burlingame, CA, USA). Images were acquired using a slide scanning system (Panoramic Scan, 3DHistech, Ltd., Budapest, Hungary).

### Statistical Analysis

#### Data acquisition and analysis with commercial systems

Data acquisition and analysis were performed using TDT processors and Opensorter (TDT, Alachua, FL, USA), Synapse Software (Synapse Software, Colorado Springs, CO, USA), and NeuroExplorer (NEX Technology, Colorado Springs, CO, USA). MATLAB (MathWorks, Natick, MA, USA) and GraphPad Prism 6.0 (GraphPad Software, San Diego, CA, USA) were used for further statistical analyses. All data are presented as mean□±□standard error, unless otherwise indicated. Statistical comparisons of two variables from an identical set of subjects were performed using paired *t*-tests. Otherwise, an unpaired *t*-test or Mann–Whitney U test was used. Statistical significance was defined as p-values <□0.05, <0.01, and <0.001, as indicated.

#### Stimulation artifact removal using the Period-based Artifact Reconstruction and Removal Method (PARRM) algorithm

As we used a threshold-based spike-sorting algorithm, we sought to remove artifacts caused by stimulation. We adopted the PARRM algorithm: the latest stimulation artifact removal method requiring only a few hyperparameters to be tuned (Dastin-van Rijn et al., 2021). We adjusted the parameters (stimRate = 140, fs = 24,414, winSize = 10·fs, skipSize = floor(fs·0.01), winDir = “both”, and perDist = 0.1) and applied the algorithm to the data for the stimulation period only.

#### Spike-sorting using the wave_clus algorithm

For the stimulation artifact removed data, we sorted single units with the wave_clus algorithm, an automatic spike-sorting algorithm based on wavelet coefficients (Chaure et al., 2018). We adjusted the parameters as follows: sr = 24,414, w_pre = 20, w_post = 44, alignment_window = 20, detection = “neg”. After the automatic sorting procedure, we manually merged and split the clusters based on the feature space extracted using the algorithm.

#### Parameterization of the aperiodic component in LFP using the SpecParam algorithm

To parameterize the aperiodic component of LFP, we used the SpecParam algorithm, also known as Fitting Oscillations and One-Over-f (FOOOF) (Donoghue et al., 2020). Before using the SpecParam algorithm, we linearly interpolated the neural power spectral densities at approximately 60 Hz in the log-log space (i.e., the logarithm of frequency and power) to relieve the notch filter effects. We adjusted the parameters (max_n_peaks = 4, min_peak_height = 0.15, peak_width_limts = [1; 20], and aperiodic_mode = “fixed”) and used the input frequency range of 6–110 Hz to avoid oscillations crossing the fitting range borders (84). Note, here we used the “fixed” mode (i.e., the knee parameter k=0), because the “knee” mode generated worse fits for several data sets than the “fixed” mode.

**Declaration of Competing Interest:** Authors declare that they have no competing interests.

## Funding

Korea Brain Research Institute Research Programs, grant 23-BR-01-01, 23-BR-03-01, and 23-BR-04-04 (JCR).

National Research Foundation of Korea, funded by the Ministry of Science and Information and Communication Technology, grant NRF-2017M3A9G8084463 and NRF-2022R1A2C1004216 (JCR).

Korea Research Institute of Bioscience and Biotechnology (KRIBB) Research Initiative Program (KGM5282322) (JCR).

We thank Drs. John Isaac, Jaewon Ko, and Jinhyun Kim for their helpful comments. Schematic diagrams were adapted from reset images of BioRender.com, and grammatical errors were corrected by Elsevier Language Editing Service, Grammarly, and ChatGPT.

## Author contributions

Study design: J-CR, ISC

Data gathering and analysis: ISC

Further analysis of the *in-vivo* electrophysiology data: JMK

Project supervision: J-CR, JWC

Writing: All authors contributed

**Data and materials availability:** All data are available at https://doi.org/10.6084/m9.figshare.25859365.v1

