## supplemental figure 10 for "Modulation of premotor cortex excitability mitigates the behavioral and electrophysiological abnormalities in a Parkinson’s Disease Mouse Model"

Jong-Cheol Rah

**This PDF file includes:**

Supporting text

Figures S1 to S10

**Fig. S1. Unilaterally lesioned 6-OHDA parkinsonian mouse model**

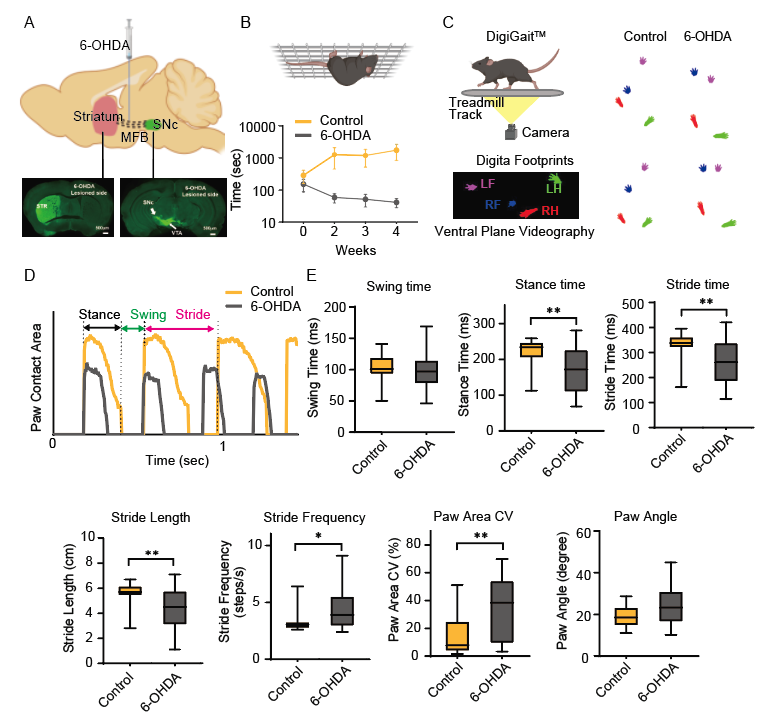

**Fig. S1.**

**A.** Stereotactic microinjection of 6-OHDA into the medial forebrain bundle (MFB) to establish a unilateral parkinsonian mouse model (top). Tyrosine hydroxylase (TH) immunohistochemistry in the striatum and substantia nigra compacta after MFB 6-OHDA lesions (bottom). STR, striatum; SNc, substantia nigra compacta; VTA, ventral tegmental area. **B.** Mean hanging time measured in the wire-hanging test after MFB 6-OHDA lesions (4 weeks after 6-OHDA-lesion, 41.18 ± 12.91 s, n = 8 mice vs. 1748.33 ± 911.44 s for sham control mice, n = 6 mice, p = 0.048, Student’s t-test). **C.** Diagram of the experimental setup to measure gait (left) and representative positions of paws with a walking speed of 17 cm/s extracted from the ventral plane videography (right). **D.** Paw areas with left hind limbs in contact with treadmill surface. **E.** Gait parameters of the contralateral hindlimb paw. Sham control (yellow) and 6-OHDA-lesioned (gray) mice. Swing time (6-OHDA-lesioned mice, 97.91 ± 4.01 ms, n = 45 mice; control mice, 103 ± 6.92 ms, n = 11 mice, p = 0.377, Mann–Whitney test), stance time (6-OHDA-lesioned mice, 165.95 ± 9.22 ms, n = 45 mice; control mice, 222 ± 12.46 ms, n = 11 mice, p = 0.003, Mann–Whitney test), stride time (6-OHDA-lesioned mice, 262.53 ± 11.8 ms, n = 45 mice; control mice, 325 ± 18.15 ms, n = 11 mice, p = 0.007, Mann–Whitney test), stride length (6-OHDA-lesioned mice, 4.41 ± 0.21 cm, n = 45 mice; control mice, 5.52 ± 0.3 cm, n = 11 mice, p = 0.008, Mann–Whitney test), stride frequency (6-OHDA-lesioned mice, 4.32 ± 0.23 steps/s, n = 45 mice; control mice, 3.30 ± 0.31 steps/s, n = 11 mice, p = 0.018, Mann–Whitney test), paw area CV (6-OHDA-lesioned mice, 34.52 ± 3.25 %, n = 45 mice; control mice, 15.93 ± 4.51%, n = 11 mice, p = 0.007, Mann–Whitney test), paw angle (6-OHDA-lesioned mice, 23.90 ± 1.21°, n = 45 mice; control mice, 19.12 ± 1.68°, n = 11 mice, p = 0.062, Mann–Whitney test). Data are presented as box and whisker plots, where each dot represents the mean of control and 6-OHDA lesioned mice.

**Fig. S2. Spike sorting of M2 recordings**

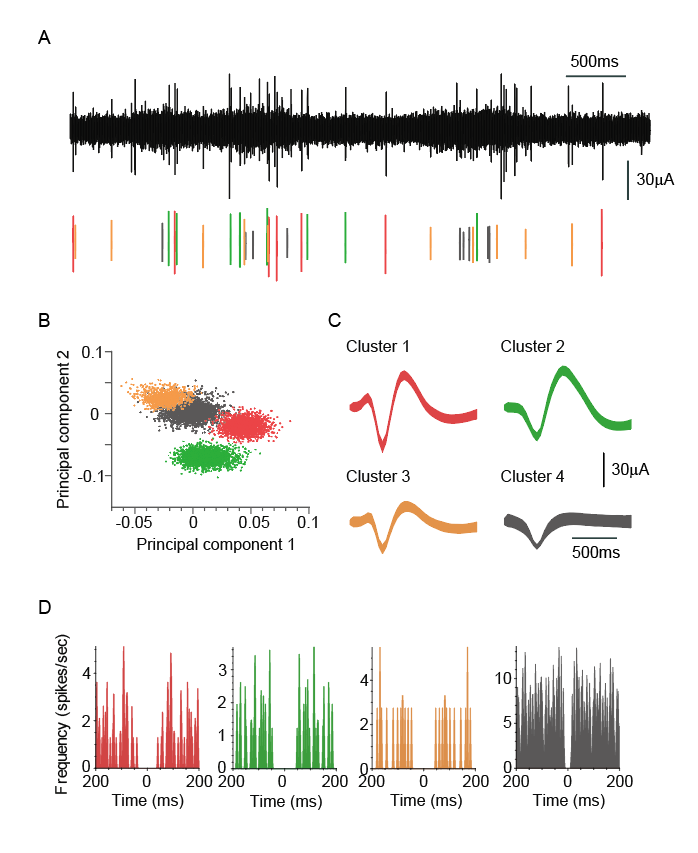

**Fig. S2.**

**A.** Lowpass filtered electrophysiological recoding from M2. Events whose amplitudes greater than a set threshold (>4x standard deviation) are marked by colored vertical lines representing the time of the events. **B.** Scatter plot of spike waveforms (red, green, or yellow) and noise cluster (gray) in the space of principal components. **C.** Three distinct single units sorted from M2 recordings and the noise cluster. **D.** Autocorrelograms of each cluster with bin width of 0.1 ms.

**Fig. S3. LFP power of M2 in the 6-OHDA-lesioned and control mice**

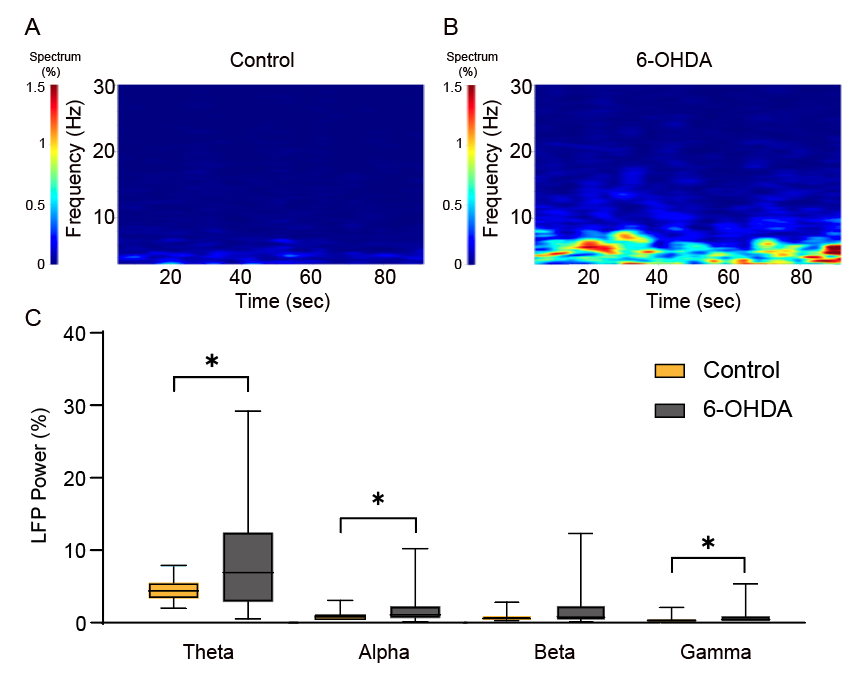

**Fig. S3.**

**A-B.** Time-frequency plots of LFP spectral power in M2 in control mice (A) and 6-OHDA lesioned mice (B). **C.** Bar graphs represent the relative LFP power in M2 in control and 6-OHDA lesioned mice. Power in the frequency ranges of theta (3–7 Hz), alpha (7–12 Hz), beta (12–30 Hz), and gamma (30–80 Hz) is normalized to the total LFP energy of the control. Control (n = 26) and (n = 81) lesioned mice. (*p < 0.05, Mann–Whitney test). Data are presented as box and whisker plots, where each dot represents the mean of control and 6-OHDA lesioned mice.

**Fig. S4. Spontaneous synaptic transmission of the controls and PD mouse model**

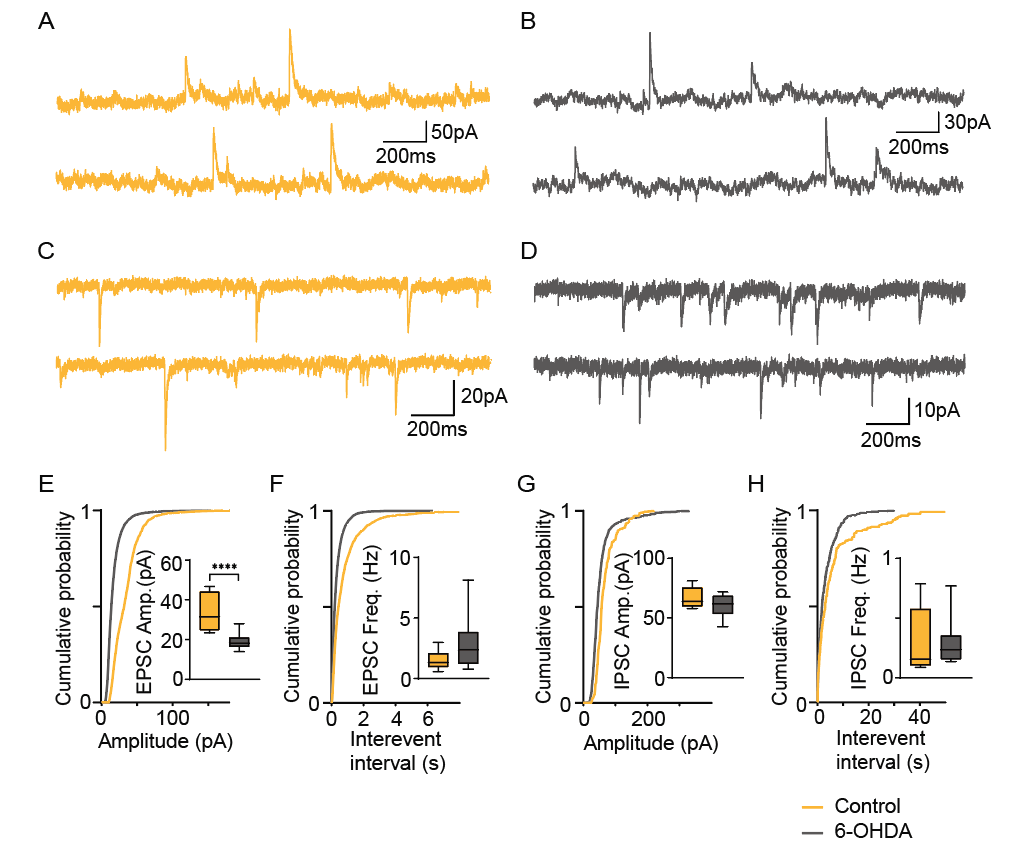

**Fig. S4.**

**A-D.** Example traces of sEPSCs (A, B) and sIPSCs (C, D) measured from the 6-OHDA-lesioned mice (gray) and control mice (yellow). sEPSCs are recorded at -70 mV, and sIPSCs were recorded at a V_H_ of 0 mV. **E-F**. The cumulative distribution function for the amplitude (E, 6-OHDA-lesioned mice, 19.30 ± 1.02 pA, n = 18 mice; control mice, 34.69 ± 3.25 pA, n = 9 mice, p < 0.0001, Mann–Whitney test) and interevent interval (F, 6-OHDA-lesioned mice, 2.86 ± 0.45 Hz, n = 18 mice; control mice, 34.69 ± 3.25 pA, n = 9 mice, p = 0.075, Mann–Whitney test) of sEPSCs recorded 6-OHDA lesioned mice (gray) and control mice (yellow). The insets show the mean and SEM from nine animals. **G-H.** Same as E and F, but for the sIPSCs. The cumulative distribution function for the amplitude (G, 6-OHDA-lesioned mice, 60.07 ± 3.42 pA, n = 8 mice; control mice, 67.03 ± 3.21 pA, n = 7 mice, p = 0.281, Mann–Whitney test) and interevent interval (H, 6-OHDA-lesioned mice, 0.30 ± 0.07 Hz, n = 8 mice; control mice, 0.29 ± 0.10 Hz, n = 7 mice, p = 0.463, Mann–Whitney test) of sIPSCs recorded 6-OHDA lesioned mice (gray) and control mice (yellow). Insets show the mean and SEM from six animals.

**Fig. S5. M2 activity upon surprise-induced licking not interruptions**

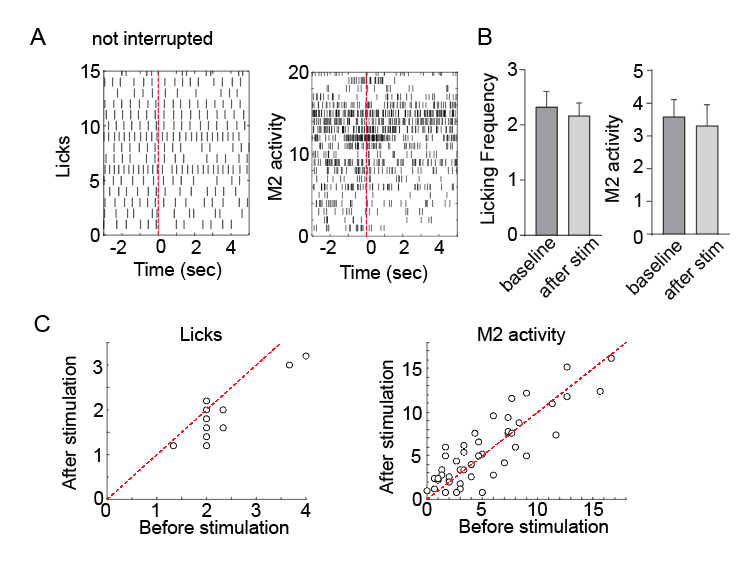

**Fig. S5.**

**A.** The raster presentation of licks and simultaneously recorded M2 activity in exemplified trials. **B.** Averaged frequencies of licking and M2 activity in the trials not interrupted by the noise (licking frequency (left), n = 9 mice; M2 activity (right), n = 9 mice). **C.** Scatter plots of licking and M2 activity frequencies before and after noise stimulation without significant interruption.

**Fig. S6. Analysis of LFP measured in the M2 of the PD mouse model with optogenetic stimulation of STN-projecting M2 neurons**

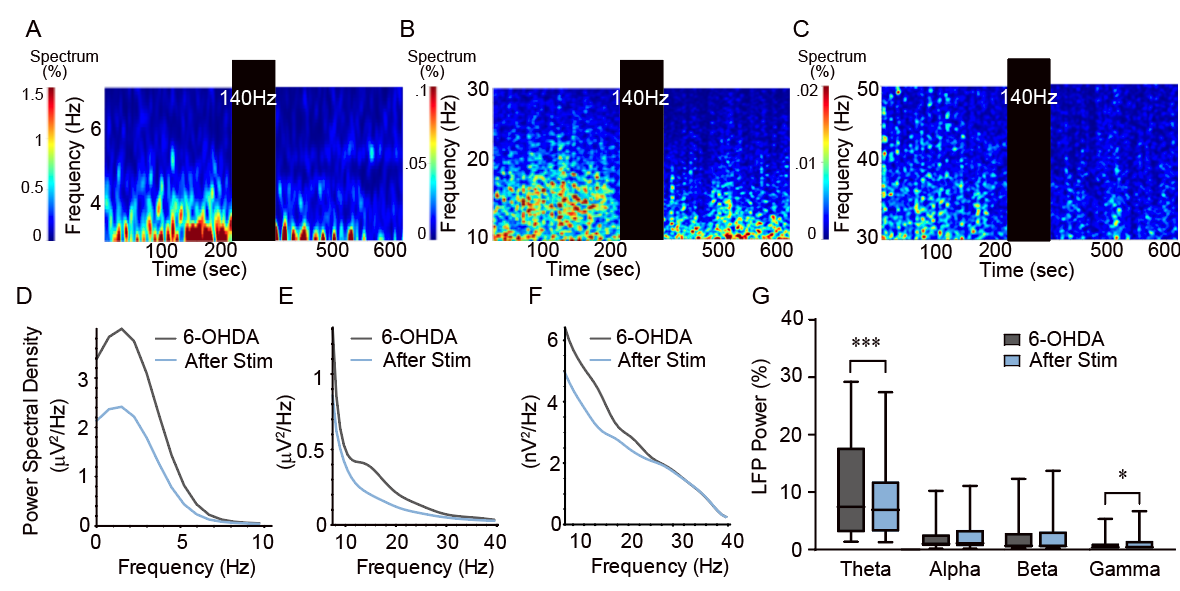

**Fig. S6.**

**A-C** Representative spectrograms of the local field potential (LFP) with theta (A), beta (B), and gamma (C) frequency ranges. STN-projecting M2 neurons are stimulated at 140 Hz for 3 min, as indicated (black boxes). **D-F.** Average LFP power at theta (D), beta (E), and gamma (F) frequency ranges before and after stimulation. **G.** Relative LFP power changes at various frequency ranges measured at the M2 by optogenetic stimulation (n = 45). The relative LFP power is calculated by normalizing the energy of the frequency range to the total energy at the baseline period. Frequency ranges are defined as theta (3–7 Hz), alpha (7–12 Hz), beta (12–30 Hz), and gamma (30–50 Hz). LFP power is significantly decreased in the theta and gamma ranges (***p<0.001, *p<0.05 relative to 6-OHDA lesioned mice; paired t-test, n = 45 neurs).

**Fig. S7. Classification of neurons based on spike waveform**

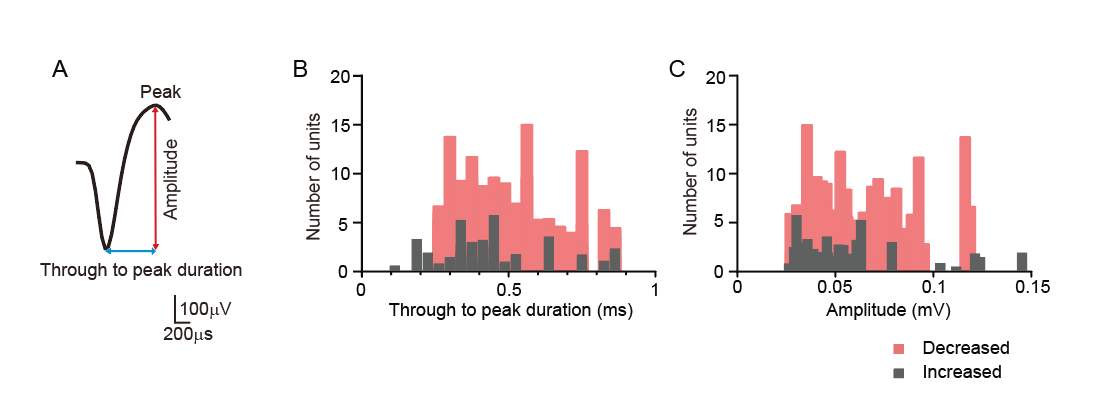

**Fig. S7.**

**A.** Example of an averaged spike waveform obtained from M2 neurons. **B-C.** Baseline frequency of isolated units as functions of kinetic properties. The histograms of units that showed increased or decreased frequencies upon optogenetic stimulation are colored in back and red respectively (increased, n = 41; decreased, n = 83).

**Fig. S8. Comparison of single unit activities from each hemispheric M2 of the hemi-Parkinsonian mouse model**

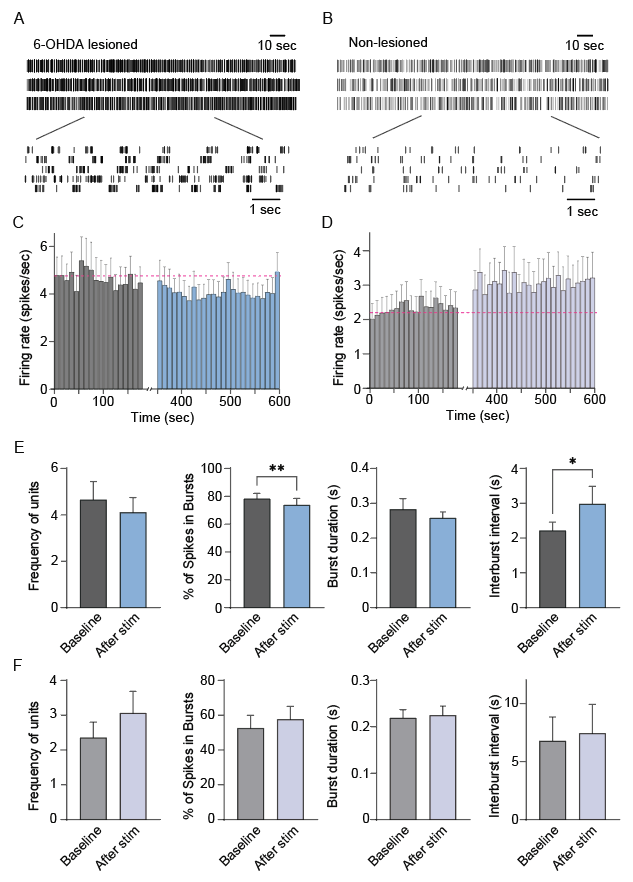

**Fig. S8.**

**A-B.** Example raster plots of representative single units with electrical stimulation in the 6-OHDA lesioned (A) and non-lesioned (B) hemispheres. **C-D.** Averaged unit counts binned for 10 s from the 6-OHDA lesioned (C) and non-lesioned (D) hemispheres of the hemi-Parkinsonian mouse model (lesioned hemispheres, n =25, 3 mice; non-lesioned hemispheres, n =19, 3 mice). **E-F**. Frequency, spikes in burst, burst duration, and interburst interval of M2 units upon high-frequency electrical stimulation (lesioned hemispheres (E), n =25, 3 mice; non-lesioned hemispheres (F), n =19, 3 mice, ^**^p < 0.01, ^*^p < 0.05, paired t-test).

**Fig. S9. Spontaneous synaptic transmission of the PD mouse model**

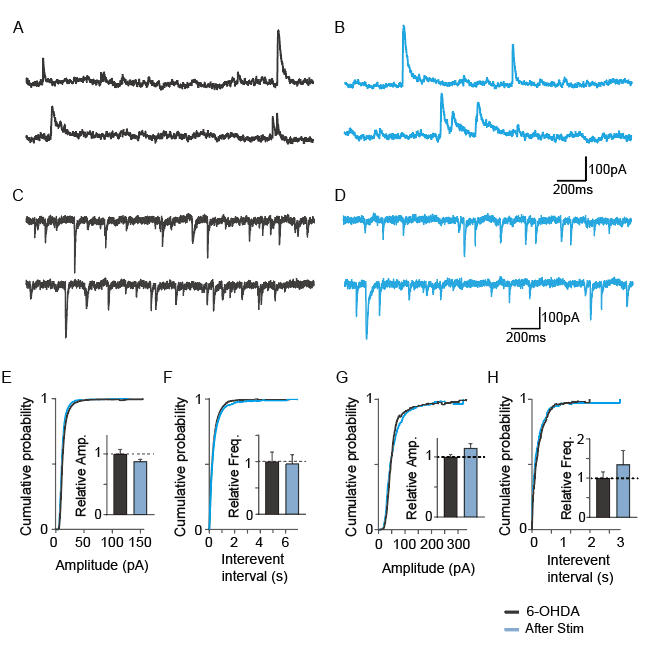

**Fig. S9.**

**A-B.** Example traces of sEPSCs (A) and sIPSCs (B) measured from the 6-OHDA lesioned PD mouse model before (gray) and after (blue) optogenetic stimulation. sEPSCs are recorded at -70 mV, and sIPSCs are recorded at 0 mV. **C-D**. The cumulative distribution function for the amplitude (C) and interevent interval (D) of sEPSCs recorded before (gray) and after stimulation (blue). The insets show the mean and SEM from nine animals. **E-F.** Same as C and D, but for the sIPSCs.

**Fig. S10. Changes in the aperiodic component of LFP in the STN according to the optogenetics stimulation.**

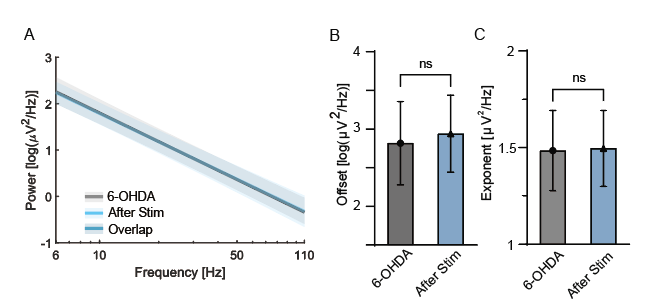

**Fig. S10.**

**A.** Comparison of aperiodic parameters before (gray) and after (blue) the stimulation. The aperiodic fit per animal is estimated by averaging the aperiodic fits obtained from STN channels. The average fit for each condition is calculated by averaging the aperiodic fits obtained from animals in each condition, as shown by the thick gray and blue lines (before stimulation: gray, after stimulation: blue). The shaded areas represent the standard deviation for each condition. **B-C.** Comparisons of aperiodic offset and exponent before and after stimulation. The left (B) and right (C) panels describe the aperiodic offset (Friedman test with Duun’s multiple comparisons test; n = 9) and the aperiodic exponent (one-way repeated measure ANOVA with Bonferroni’s multiple comparisons test; n = 9, *𝑝<0.05), respectively.
